# The persistent effects of predator odor stressor enhance interoceptive sensitivity to alcohol through GABA_A_ receptor adaptations in the prelimbic cortex in male, but not female rats

**DOI:** 10.1101/2024.10.30.621141

**Authors:** Ryan E. Tyler, Maya N. Bluitt, Kalynn J. Van Voorhies, Wen Liu, Sarah N. Magee, Elisabeth R. Pitrolo, Victoria L. Cordero, Laura C. Ornelas, Caroline G. Krieman, Brooke N. Bender, Alejandro M. Mosera, Joyce Besheer

**Affiliations:** Neuroscience Curriculum, School of Medicine, University of North Carolina – Chapel Hill, Chapel Hill, NC; Bowles Center for Alcohol Studies, School of Medicine, University of North Carolina, Chapel Hill, NC; Department of Psychiatry, School of Medicine, University of North Carolina, Chapel Hill, NC

**Author notes:** Correspondence: Joyce Besheer, Ph.D. Professor Bowles Center for Alcohol Studies Department of Psychiatry University of North Carolina at Chapel Hill Chapel Hill, NC 27599-7171.

**Keywords:** drug discrimination, anterior insular cortex, 2, 5-dihydro-2, 4, 5-trimethylthiazoline, individual differences, PTSD, alcohol use disorder

## Abstract

**Background:** Traumatic stress is associated with high rates of problematic alcohol use, but how the persistent effects of trauma impact sensitivity to alcohol remain unknown. This study examined the persistent effects of traumatic stress exposure on sensitivity to alcohol and underlying neurobiological mechanisms in rats.

**Methods:** Male (N=98) and female (N=98) Long-Evans rats were exposed to the predator odor TMT, and two weeks later, molecular, neuronal, and behavioral sensitivity to alcohol were assessed. Next, rats were trained to discriminate alcohol from water (male N=70; female N=56), and the impact of TMT on interoceptive sensitivity to alcohol and the alcohol-like effects of systemic GABA_A_ receptor activation were evaluated. Lastly, functional involvement of GABA_A_ and NMDA receptors in the prelimbic cortex (PrL) and the anterior insular cortex (aIC) was investigated.

**Results:** TMT exposure sex-dependently altered PrL *Gabra1*, and elevated aIC *Grin2b* and *Grin2c* in males. TMT increased PrL c-Fos in males, which was attenuated by alcohol administration. Alcohol-induced locomotor and startle response effects were attenuated in the TMT group in both sexes. TMT exposure potentiated interoceptive sensitivity to alcohol in males but not in females, and this effect was driven by GABA_A_ receptors in the PrL. Greater stress reactivity during TMT exposure was associated with higher interoceptive sensitivity to alcohol, and alcohol exposure history was linked to a heightened stress response to TMT in males.

**Conclusions:** Traumatic stress increased interoceptive sensitivity to alcohol in males, but not females, through PrL GABA_A_ receptor adaptations, potentially enhancing the stimulatory, and by extension the rewarding, effects of alcohol.

## INTRODUCTION

Post-traumatic stress disorder (PTSD) is a debilitating neuropsychiatric condition that develops in some individuals following exposure to a traumatic event(1). PTSD is highly co- morbid with alcohol use disorder (AUD)(2–4), and stress is commonly linked to increased alcohol consumption or craving across multiple species(5–13). Sensitivity to the subjective effects of alcohol is a strong predictor of AUD development and associated with differences in alcohol use(14–16). However, how the persistent effects of trauma impact sensitivity to alcohol, and associated neurobiological mechanisms, are unknown.

In animal models, sensitivity to the effects of alcohol can be assessed using drug discrimination procedures(17–35). The learned behavior is guided by the internal state of the animal following alcohol administration, and as such we will refer to these alcohol effects as interoceptive effects(17, 24–26, 36, 37). There is an extensive literature to support that the interoceptive effects of alcohol are predominantly driven by the effects of alcohol on GABA_A_ and NMDA receptors(19, 30, 31, 38–40). Generally, GABA_A_ receptor-mediated disinhibition by alcohol is associated with the stimulatory effects of alcohol(41–44), and NMDA receptor inhibition by alcohol is associated with the sedative effects(43, 45, 46). Furthermore, stress-induced neuroadaptations are driven in part by synaptic plasticity induced by adaptations in NMDA and GABA_A_ receptors(47–53), suggesting potential interactions and overlapping mechanisms between stress and the expression of the interoceptive effects of alcohol.

Furthermore, the prelimbic cortex (PrL) and the anterior insular cortex (aIC) are highly sensitive to acute and persistent effects of stress(48, 54, 55), as well as having involvement in the subjective / interoceptive effects of alcohol(19, 20, 22, 37, 40). Additionally, both brain regions are strongly implicated in PTSD and AUD(56–66). As such, we hypothesized that traumatic stress may impact interoceptive sensitivity to alcohol through changes in the GABA_A_ and/or NMDA receptor in the PrL and/or the aIC.

Exposure to a predator odor has been used as an animal model of a traumatic stressor, and the resulting behavioral and neural adaptations have some relevance to understanding symptoms associated with PTSD(5–7, 67–81). In the present work, rats were exposed to the scent of a predator (2,5-dihydro-2,4,5-trimethylthiazoline (TMT), a synthetic component in fox feces) in an inescapable enclosure to model a traumatic stressor(5, 6, 67–70, 72, 73, 79, 81–84). Outcomes were always collected two weeks after the TMT exposure to assess persistent effects which more closely models the lasting effects of trauma that is observed in people with PTSD. We recently showed that TMT exposure potentiated the interoceptive effects of alcohol in a small sample of male rats using an operant drug discrimination procedure(24). The present work expands on this finding using Pavlovian drug discrimination methods in both sexes and further investigates the neurobiological and behavioral mechanisms that underlie this effect.

We began by assessing the effects of TMT stressor exposure on GABA_A_ and NMDA receptor subunit gene expression in the PrL and aIC. Next, we evaluated the impact of TMT exposure and acute alcohol administration on c-Fos immunoreactivity in the PrL and aIC, locomotor activity, and startle response behavior. Following this, we investigated the effect of TMT exposure on interoceptive sensitivity to alcohol using a Pavlovian drug discrimination training procedure. Additionally, we examined the functional role of GABA_A_ and NMDA receptor adaptations in the PrL and aIC through pharmacological approaches. Finally, we investigated how individual differences in stress reactivity during TMT exposure was associated with the interoceptive effects of alcohol. This is an important consideration given that only a minority of individuals who experience trauma develop PTSD(1), and an even smaller percent will develop a comorbid AUD. Together, these experiments may help us better understand the high co-morbidity between PTSD and AUD by elucidating how traumatic stress affects sensitivity to alcohol.

## METHODS AND MATERIALS

Male and female Long-Evans rats were exposed to TMT for 15-min and all outcome measures were assessed two weeks later (rats remained undisturbed in their home cage during the two week “incubation” period). Neurobiological assessments focused on GABA_A_ and NMDA receptors in the PrL and aIC. Experiment 1 (gene expression, Fig. 1, Supplemental Table 1 shows primer sequences), Experiment 2 (c-Fos, Fig. 2), and Experiment 3 (behavior, Fig. 3) used alcohol-naïve rats, whereas Experiment 4 (drug discrimination, Fig. 4-6) used rats trained on a Pavlovian drug discrimination procedure to discriminate the interoceptive effects of alcohol (2.0 g/kg, i.g.) from water. Bilateral cannulae were implanted into a subset of male rats targeting the prelimbic cortex (PrL) or the anterior insular cortex (aIC) based on(85). Analysis of variance (ANOVA) was run with TMT exposure as a between-subjects factor, and with alcohol as a between or within-subjects factor as appropriate. Sex was used as a between subjects factor when appropriate. Detailed methods are included in the Supplementary Materials.

**Figure 1.**
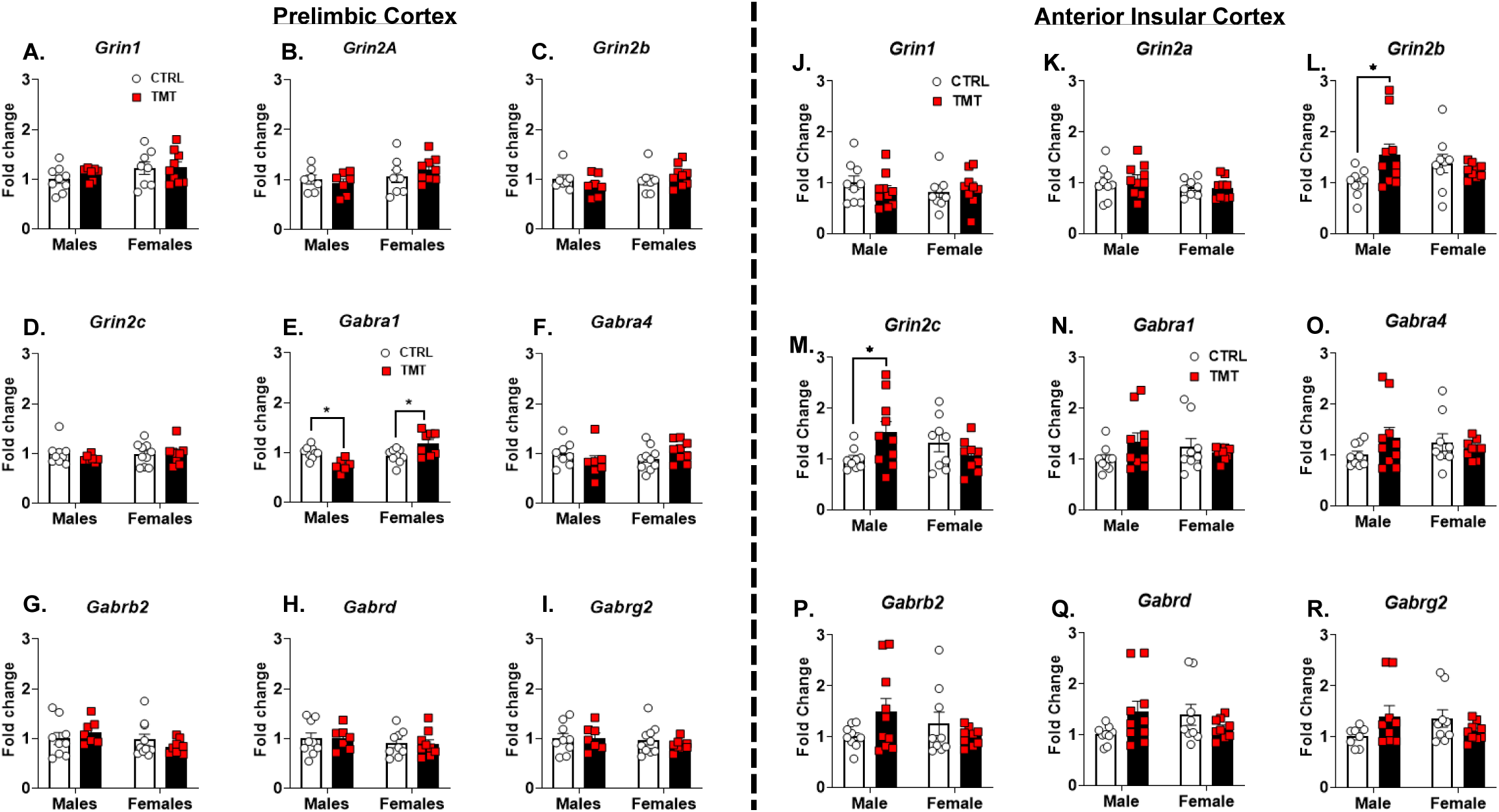
**TMT exposure sex-dependently altered *Gabra1* expression in PrL and upregulated NMDA receptor subunits in aIC in males**. For NMDA receptor sub-unit gene expression in the PrL (A-D), there was no effect of TMT exposure or sex. For the GABA_A_ α-1 sub-unit gene expression (E), TMT exposure decreased expression in males but increased expression in females (interaction: F(1, 28)=18.12, p=0.0002). Other GABA_A_ receptor sub-unit genes (F-I) were not affected by TMT exposure or sex. For NMDA receptor sub-unit gene expression in the aIC, there was no effect of TMT exposure or sex on (A) *Grin1* and (B) *Grin2a* levels. TMT exposure increased levels of (L) *Grin2b* and (M) *Grin2c* in males, but not in females (interaction: F(1, 33)=5.10, p=0.03). GABA_A_ α-1 subunit gene expression (N-R) was not affected by TMT exposure or sex. Male/CTRL n=7-9, Male/TMT n=6-10, Female/CTRL n=7-10, Female/TMT n=8-9. *p≤0.05.

**Figure 2.**
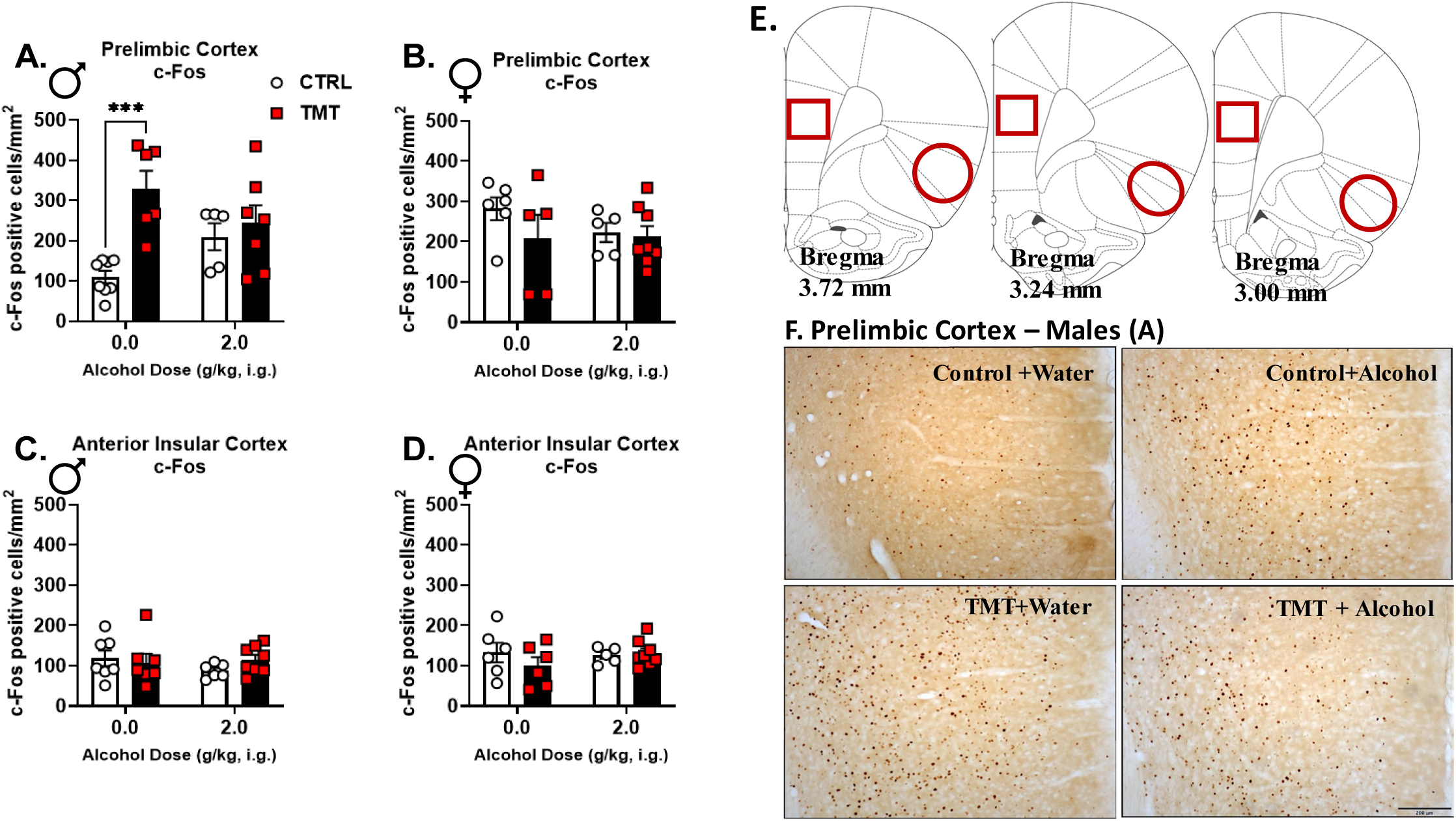
TMT exposure increased PrL c-Fos expression in males, which was not observed in rats pre-treated with alcohol. For c-Fos positive cells/mm^2^ in the PrL in males (A), TMT exposure induced an increase but alcohol did not affect c-Fos expression (TMT: F(1, 22)=12.70, p=0.002; TMT x alcohol: F(1, 22)=6.83, p=0.02). In the PrL in females (B), TMT exposure nor alcohol affected c-Fos positive cell number/mm^2^. In the aIC in males (C) and females (D), TMT exposure nor alcohol administration affected c-Fos expression. (E) Regions of quantification for the PrL (square) and the aIC (oval). (F) Representative images of Fig. 2A data – c-Fos in PrL of male rats. Scale bar is 200 μm. Male/CTRL n=6-8, Male/TMT n=6-7, Female/CTRL n=5-6, Female/TMT n=5-8. ***p≤0.001.

**Figure 3.**
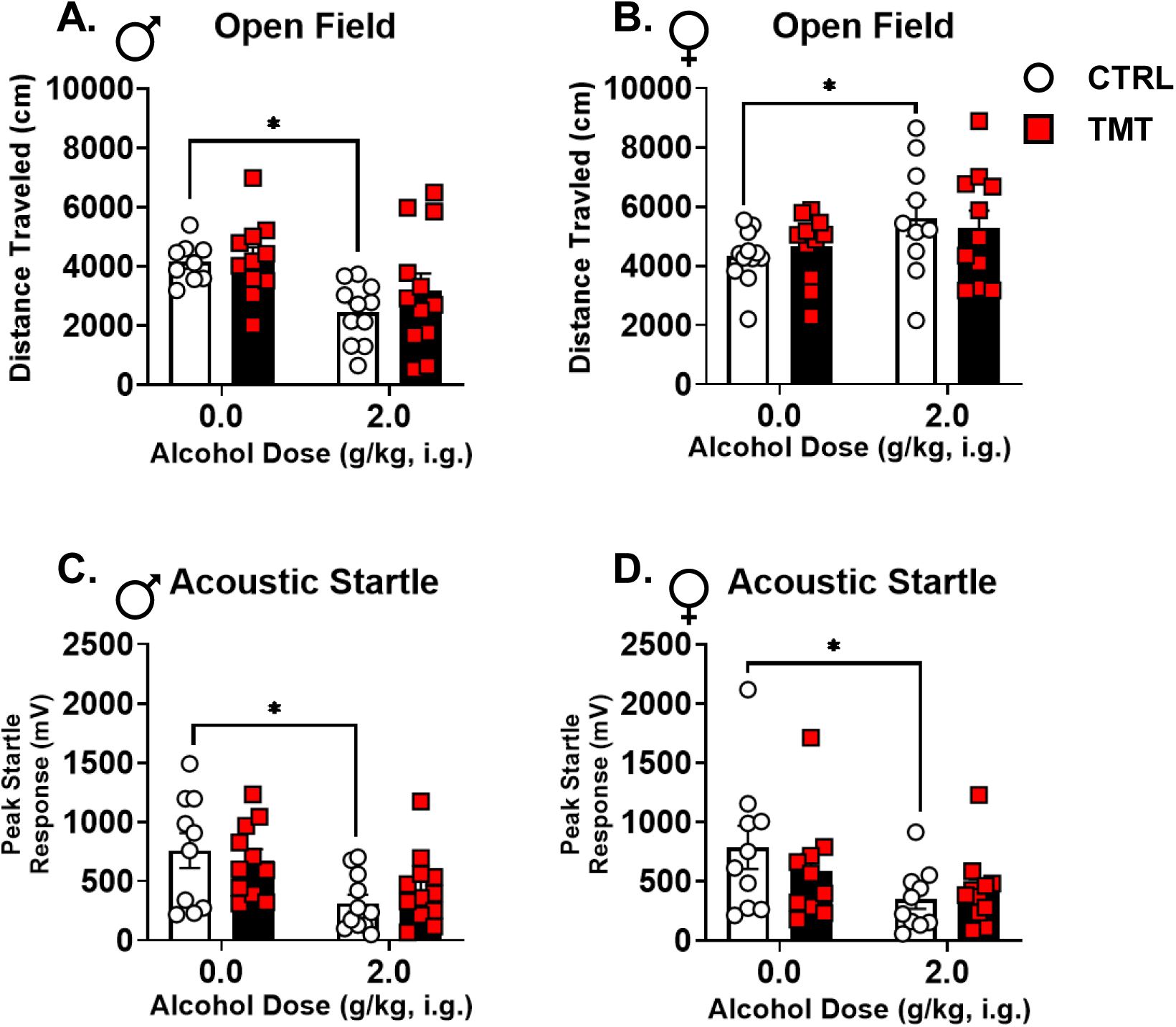
Alcohol produced locomotor and startle response effects in the control group, but not in the TMT-exposed group. For the open field test in males (A), alcohol administration (alcohol: F(1, 39)=10.63, p=0.002) decreased distance traveled in the control group (t(18)=4.31, p=0.0004), but not in the TMT group. In the open field test in females (B), alcohol administration (alcohol: F(1, 41)=4.64, p=0.04) resulted in increased distance traveled in the control group (t(20)=2.08, p=0.05), but not in the TMT group. For the acoustic startle response in males (C), alcohol administration (alcohol: F(1, 40)=11.65, p=0.002) decreased the average peak startle response in the control group (t(19)=2.79, p=0.01), but not in the TMT group. For the acoustic startle response in females (D), alcohol administration (alcohol: F(1, 37)=5.35, p=0.03) resulted in decreased average peak startle response in controls (t(18)=2.18, p=0.04), but not in the TMT group. Male/CTRL n=9-11, Male/TMT n=11-12, Female/CTRL n=10-12, Female/TMT n=10-11. *p≤0.05.

**Figure 4.**
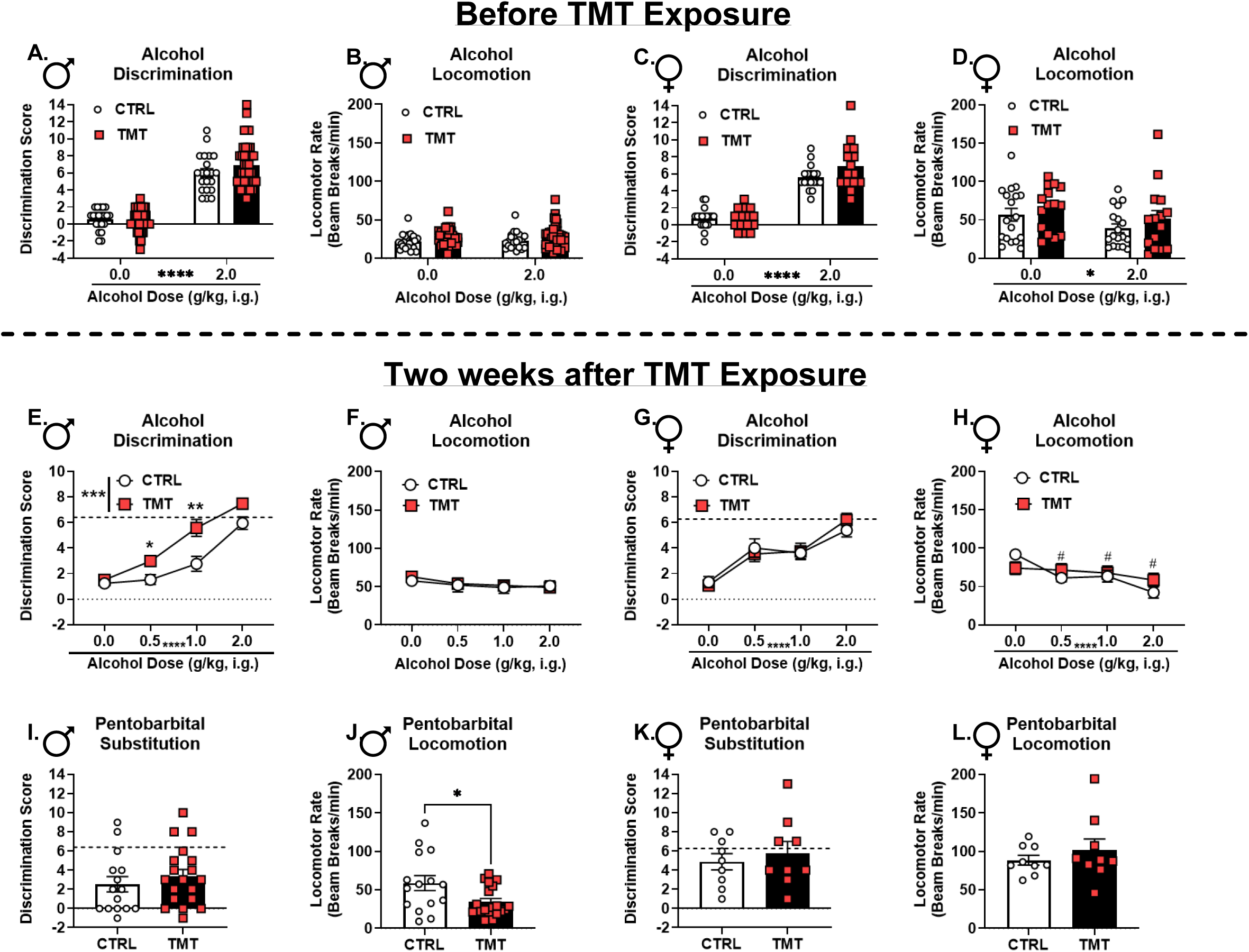
TMT exposure potentiated interoceptive sensitivity to alcohol in male but not female rats. Prior to TMT exposure, in males (A), there was a main effect of alcohol on discrimination scores (F(1, 59)=262.2, p<0.0001) as would be expected confirming that the interoceptive effects of alcohol were guiding behavior, and no difference between the TMT and control group. There was no effect on locomotor rate for alcohol administration or group (TMT vs. CTRL). Prior to TMT exposure in females, there was a main effect of alcohol on discrimination scores (F(1, 31)=211.8, p<0.0001) and no difference between groups. Likewise for locomotor rate (D), there was a main effect of alcohol (F(1, 31)=6.48, p=0.02), but no group differences. Following a 2-week period after the TMT exposure, (E) male rats exposed to TMT showed potentiated sensitivity to the interoceptive effects of alcohol (TMT: F(1, 59)=11.32, p=0.001; alcohol dose: F(2.38, 140.3)=63.83, p<0.0001; TMT x alcohol dose: F (3, 177) = 3.09, p = 0.03) specifically at the 0.5 (p=0.02) and 1.0 g/kg (p=0.009) alcohol doses. (F) There was no effect of TMT exposure or alcohol dose on locomotor rate. In females after TMT exposure, (G) there was a main effect of alcohol dose (F(3, 93)=27.03, p<0.0001), but no effect of TMT exposure on discrimination scores. For locomotion in females, (H) there was a main effect of alcohol dose (F(3, 93)=11.77, p<0.0001), no effect of TMT exposure, but there was a significant alcohol dose by TMT exposure interaction (F(3, 93)=3.70, p=0.01). Locomotion was decreased at every alcohol dose compared to vehicle (^#^p≤0.05) in controls, but not in the TMT group. For pentobarbital substitution in males, (I) there was no effect of TMT exposure on discrimination scores, but (J) the TMT group showed a decreased locomotor rate compared to controls (t(32)=2.45, p=0.02). Pentobarbital did not produce substitution for the alcohol training dose in males. For pentobarbital substitution in females, there was no effect of TMT exposure on (K) discrimination scores or (L) locomotor rate. Pentobarbital produced substitution for the alcohol training dose in females. The dotted lines on panels E, G, I, and K represent the mean (groups combined because this was prior to the TMT exposure) of the alcohol session discrimination scores prior to the TMT exposure taken from panel A and C, receptively, and serve as a comparator for full substitution of the 2 g/kg alcohol training dose. For males, the mean was 6.41 ± 0.42 SEM, and for females the mean was 6.27 ± 0.23 SEM. Discrimination scores reflect the degree of substitution for the 2.0 g/kg alcohol training dose, or the 2.0 g/kg alcohol-like effects. Alcohol dose response (A-H) sample sizes: Male/CTRL n=21, Male/TMT n=40, Female/CTRL n=18, Female/TMT n=15. Pentobarbital substitution (I-L) sample sizes: Male/CTRL n=15, Male/TMT n=19, Female/CTRL n=9, Female/TMT n=9. Only rats that met criteria for discrimination before and after the TMT exposure were included in analyses. Discrimination was defined as an alcohol session discrimination score ≥ 2 the water session score. *p≤0.05. **p≤0.01. ****p≤0.0001. ^#^p≤0.05 relative to vehicle in the CTRL group.

## RESULTS

### TMT exposure sex-dependently altered Gabra1 expression in PrL and upregulated NMDA receptor subunits in aIC in males (***Figure 1***)

Given that the interoceptive effects of alcohol are primarily modulated by GABA_A_ and NMDA receptors(19, 20, 30, 31, 38–40), we initially investigated the impact of TMT exposure on GABA_A_ and NMDA receptor subunit gene expression two weeks post-exposure in the prelimbic cortex (PrL) and anterior insular cortex (aIC). In the PrL, TMT exposure did not affect NMDA receptor gene expression (Fig. 1A-D). GABA_A_ subunit expression was mostly unaffected by TMT except for *Gabra1*, which was decreased in males and increased in females (Fig. 1E). In the aIC, the NMDA subunits *Grin2b* and *Grin2c* were upregulated in males, but not in females (Fig. 1L, M). No changes in the GABA_A_ subunits were observed in the aIC (Fig. 1N-R). These results indicate that TMT exposure affects GABA_A_ receptor gene expression in the PrL in a sex-dependent manner and increases NMDA receptor gene expression in the aIC in males.

### TMT exposure increased PrL c-Fos expression in males, which was not observed in rats pre-treated with alcohol (***Figure 2***)

The observed changes in NMDA and GABA_A_ receptor subunit expression following TMT exposure suggested a potential alteration in the neuronal response to alcohol. Therefore, we examined neuronal activation, measured by c-Fos immunoreactivity (IR), following acute alcohol administration (0 or 2 g/kg) two weeks after TMT exposure. Males exposed to TMT showed increased c-Fos IR in the PrL following water (vehicle) administration (Fig. 2A), demonstrating increased neuronal activation under vehicle conditions. This increase was not observed in the alcohol-treated group. In contrast, no TMT or alcohol effects were observed in the females (Fig. 2B) and no changes were observed in the aIC in males or females (Fig. 2C, D). These findings indicate a lasting increase in neuronal activation following TMT exposure in the PrL in males that was attenuated by acute alcohol pretreatment, which may indicate that this stressor-induced adaptation can be remediated by alcohol use. The increase in neuronal activity in the PrL may be related to the decrease in *Gabra1* observed in the same brain region (Fig. 1).

### Alcohol produced locomotor and startle response effects in the control group, but not in the TMT-exposed group (***Figure 3***)

Given that TMT exposure altered gene expression and c-Fos IR in the brain, we next evaluated whether TMT exposure would affect behavioral sensitivity to alcohol. In the open field, alcohol (2 g/kg) decreased locomotion in males that was driven by a reduction in the alcohol-treated controls, but not the TMT group (Fig. 3A). Conversely, in the females, alcohol induced an increase in locomotor activity that was again driven by the alcohol-treated controls and not observed in the TMT group (Fig. 3B). In the ASR test, both males and females showed an alcohol-induced reduction in startle response in the control group, but not in the TMT group (Fig. 3C, D). These results indicate that TMT exposure may attenuate the locomotor and startle-response effects of alcohol.

### TMT exposure potentiated interoceptive sensitivity to alcohol in male but not female rats (***Figure 4***)

The effects of TMT exposure on (1) *Gabra1* and NMDA subunit mRNA expression in the PrL and aIC, respectively, (2) c-Fos expression in the PrL, and (3) behavioral sensitivity to alcohol suggest that TMT exposure may be altering the interoceptive effects of alcohol. As such, rats were trained to discriminate the interoceptive effects of alcohol (2 g/kg) from water. Once the discrimination was acquired (Fig. 4A, C), rats underwent TMT exposure. Sensitivity to a dose range of alcohol (0, 0.5, 1.0, 2.0 g/kg, i.g.) was tested 2 weeks later. Males that had experienced TMT exposure showed potentiated sensitivity to alcohol, specifically at lower alcohol doses (0.5 and 1.0 g/kg; Fig. 4E), with no change in general locomotion (Fig. 4F). In contrast, TMT did not alter sensitivity to the interoceptive effects of alcohol in females (Fig. 4G). In females, locomotor activity was decreased by alcohol, and the controls showed a decrease at every dose relative to vehicle, but this was not observed in the TMT group (Fig. 4H), demonstrating diminished sensitivity to the motor-suppressing effects of alcohol.

Next, we investigated the functional involvement of GABA_A_ receptors in the stressor-induced effects on alcohol interoceptive sensitivity. Pentobarbital is a GABA_A_ receptor agonist that substitutes for the interoceptive effects of alcohol(86–88). As such, the same discrimination-trained rats were administered a low dose of pentobarbital (5 mg/kg) and tested for substitution. TMT exposure did not affect the alcohol-like effects of pentobarbital in males, and there was no substitution for alcohol in males (Fig. 4I, substitution analysis relative to dotted-line).

Interestingly, the TMT group showed a lower locomotor rate during the pentobarbital test session compared to controls (Fig. 4J). TMT exposure did not alter sensitivity to the alcohol-like effects of pentobarbital or locomotor rate in females (Fig. 4K, L). However, unlike in males, females systemically administered pentobarbital showed substitution for the alcohol training dose (Fig. 4K, relative to dotted line). These data demonstrated that both males and females retained the ability to discriminate alcohol from water following TMT exposure and the 2-week incubation period. Furthermore, in males, but not females, TMT exposure potentiated interoceptive sensitivity to alcohol. TMT exposure blunted the motor-suppressing effects of alcohol in females. Pentobarbital (5 mg/kg, i.p.) produced substitution for the 2.0 g/kg training dose in female but not in male rats, and did not differ between control and TMT groups in both sexes. However, male TMT-exposed rats showed decreased locomotion after pentobarbital administration compared to control rats.

### TMT exposure potentiated the alcohol-like stimulus effects of GABA_A_ receptor agonism in the prelimbic cortex in male rats (***Figure 5***)

We next sought to identify the neurobiological mechanisms underlying the stressor-induced potentiation to the interoceptive effects of alcohol in male rats. While TMT exposure did not impact the alcohol-like interoceptive effects of the GABA_A_ agonist pentobarbital when administered systemically, given the GABA_A_ receptor adaptations in the PrL and the NMDA receptor adaptations in the aIC, we sought to assess the functional contribution of these receptors in the potentiated interoceptive sensitivity to alcohol following TMT. GABA_A_ receptor agonism and NMDA receptor antagonism produce somewhat distinct interoceptive stimuli that compose different interoceptive effects of alcohol(31).

**Figure 5.**
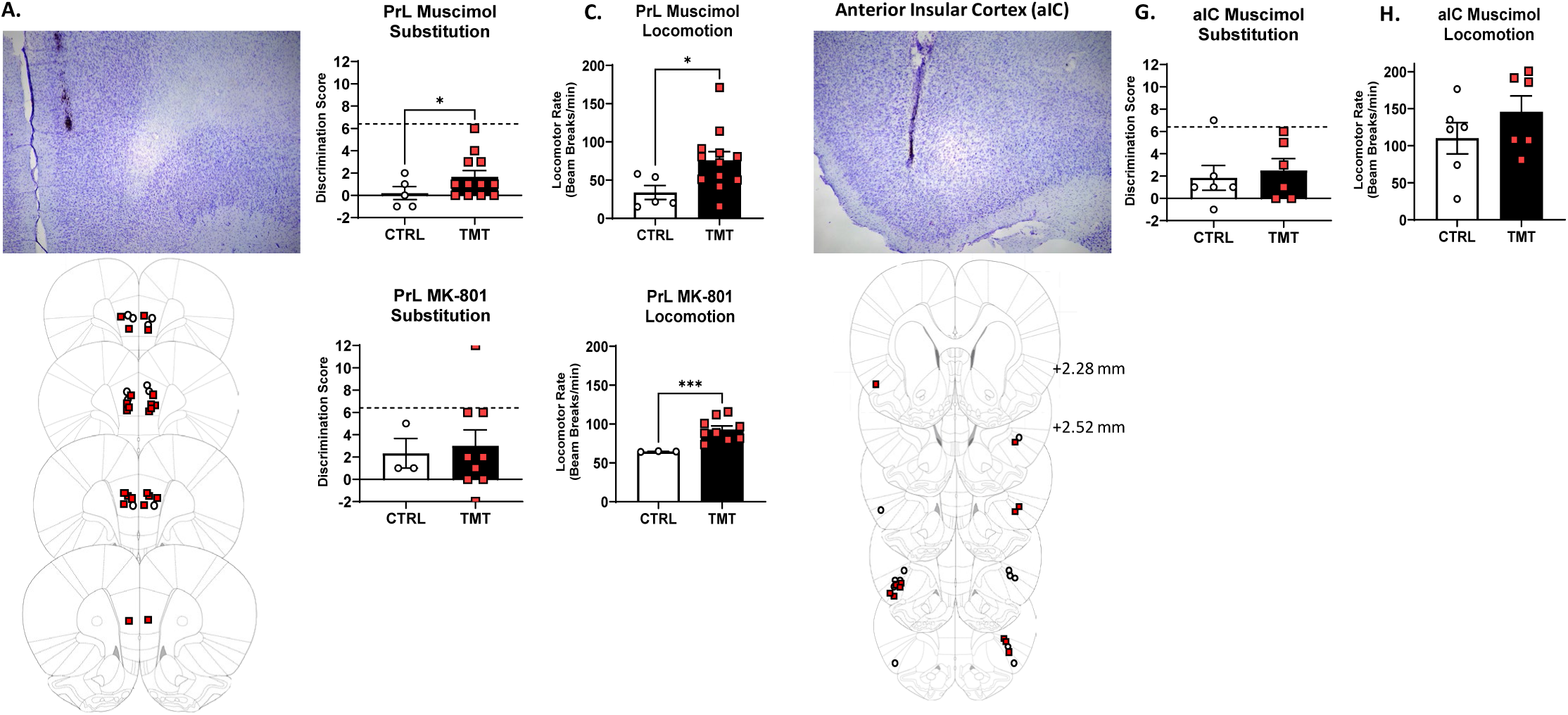
**TMT exposure potentiated the alcohol-like stimulus effects of GABA_A_ receptor agonism in the prelimbic cortex in male rats**. (A) Representative image and injector tip placement from individual rats with accurate placements in the prelimbic cortex (PrL). Site-specific injection of muscimol into the PrL of male rats resulted in (B) greater discrimination scores (t(9.05)=2.10, p=0.03) and (C) greater locomotor rate (t(14.33)=2.34, p=0.02) in the TMT group compared to controls. Site-specific injection of MK-801 into the PrL did not produce group differences for (D) discrimination scores, but did show potentiation for (E) locomotor rate (t(8.06)=5.86, p=0.0002) in the TMT group compared to controls. (F) Representative image and injector tip placement from individual rats with accurate placements in the anterior insular cortex (aIC). Site-specific injection of muscimol into the aIC of male rats resulted in no group difference for (G) discrimination scores or (H) locomotor rate. Discrimination scores reflect the degree of substitution for the 2.0 g/kg alcohol training dose, or the 2.0 g/kg alcohol-like effects. PrL/Muscimol: CTRL n=5, TMT=12. PrL/MK-801: CTRL n=3, TMT n=9. aIC/Muscimol: CTRL n=6, TMT n=6. 5 rats from the PrL and 1 from the aIC were removed from analyses due to incorrect bilateral cannulae placements. *p≤0.05. ***p≤0.001.

**Figure 6.**
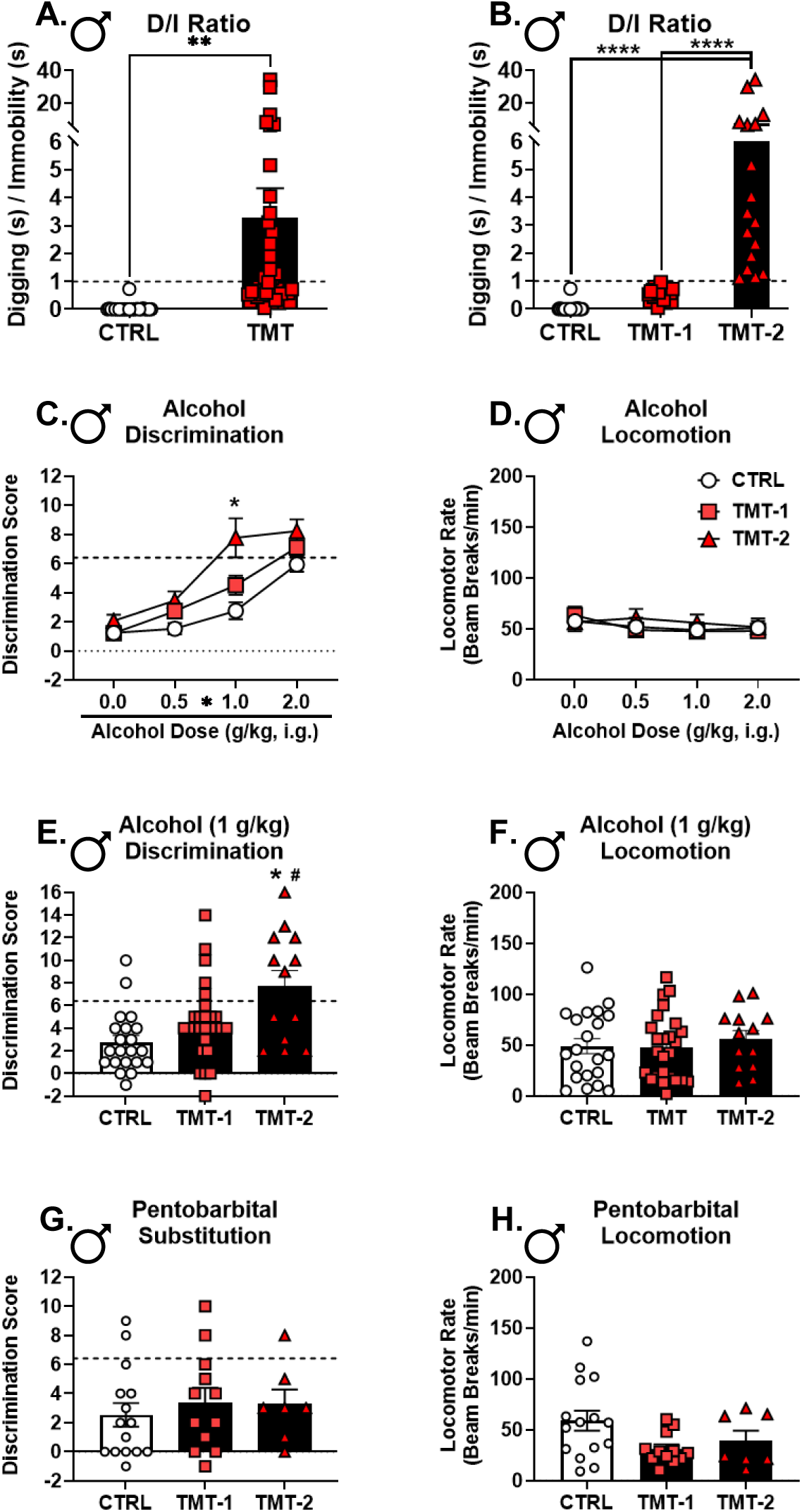
**Heightened stress reactivity to TMT exposure is associated with higher interoceptive sensitivity to alcohol, but not with the alcohol-like effects of pentobarbital**. (A) The ratio of time spent digging over time spent immobile during the 15-min TMT exposure was calculated as the D/I ratio. (B) Rats with a D/I ratio < 1 were assigned to the TMT-1 subgroup, and rats with a D/I ratio > 1 were assigned to the TMT-2 subgroup. (C) The post-TMT alcohol dose response curve was reanalyzed using the TMT subgroup. There was a main effect of group (F(2, 58)=10.20, p=0.0002), alcohol dose (F(2.43, 140.9)=74.30, p<0.0001), and group x alcohol dose interaction (F(6, 174)=2.90, p=0.01). (C) The TMT-2 group showed higher discrimination scores compared to the control group at the 1.0 g/kg alcohol dose. (D) There was no effect of alcohol dose or group for locomotor rate. (E) At the 1.0 g/kg alcohol dose, the TMT-2 group showed higher discrimination scores compared to both TMT-1 and CTRL groups (group: F(2, 58)=8.05, p=0.0008), but there were no group differences in locomotor rate (F). There were no group differences for pentobarbital (G) discrimination scores or (F) locomotor rate. Discrimination scores reflect the degree of substitution for the 2.0 g/kg alcohol training dose, or the 2.0 g/kg alcohol-like effects. Alcohol dose response sample size: CTRL n=21, TMT-1 n=27, TMT-2 n=13. Pentobarbital sample size: CTRL n=15, TMT-1 n=12, TMT-2 n= 7. *p≤0.05, **p≤0.01, ****p≤0.0001 for TMT-2 compared to controls. ^#^p≤0.05 for TMT-2 compared to TMT-1.

When the GABA_A_ agonist muscimol was injected into the PrL, the TMT group showed higher discrimination scores and locomotion compared to the control group (Fig. 5B, C). For MK-801 (NMDA receptor antagonist) injected in the PrL (Fig. 5D, E), there were no group differences in discrimination scores, but the TMT group had a higher locomotor rate during the test session. In contrast, muscimol in the aIC (Fig. 5G, H) did not produce group differences in discrimination scores or locomotion. MK-801 was not tested in the aIC. These findings indicate that the alcohol-like stimulus effects of GABA_A_ agonism in the PrL were potentiated by TMT exposure. This finding, along with gene expression and c-Fos adaptations in the PrL, indicate that GABA_A_ receptor-mediated adaptations in the PrL underlie the stressor-induced potentiation to the interoceptive effects of alcohol. Furthermore, stressor-induced potentiation in the locomotor effects following PrL muscimol or MK-801 administration suggest greater sensitivity to the stimulatory effects of alcohol.

### Heightened stress reactivity to TMT exposure was associated with higher interoceptive sensitivity to alcohol, but not with the alcohol-like effects of pentobarbital (***Figure 6***)

Having identified a brain mechanism by which TMT exposure may enhance interoceptive sensitivity to alcohol in males, we examined how individual differences in stress reactivity during the TMT exposure may be associated with the effects of TMT exposure on interoceptive sensitivity to alcohol. As such, the ratio of time spent digging over time spent immobile (D/I ratio) during the TMT exposure was used to sub-group rats into high (D/I > 1, TMT-2) and low (D/I < 1, TMT-1) stress reactivity groups (Fig. 6A, B) based on prior work from the lab(6, 67). The TMT-2 group demonstrated the highest discrimination scores at 1.0 g/kg alcohol with no effect on locomotion (Fig. 6C-F). There was no effect of TMT subgroup on the alcohol-like effects of pentobarbital or locomotion (Fig. 6G, H). These findings indicate that greater stress reactivity during the TMT exposure was associated with the greatest increase in interoceptive sensitivity to alcohol in males.

### TMT exposure increased digging and immobility behavior in male and female rats: comparing stress-reactivity in alcohol-experienced and alcohol-naïve cohorts (***Figure 7***)

Finally, to examine whether alcohol exposure history affected stress reactivity, we combined all the TMT exposure behavioral response data (behavior during the 15-min exposure) from the alcohol-naïve (Exp. 1-3) and alcohol-experienced (Exp. 4) cohorts. The alcohol-experienced rats had on average 21 days receiving 2.0 g/kg alcohol. Results indicated that male and female rats engaged in a dynamic behavioral response during the TMT exposure: digging behavior that transitioned to immobility over the course of the 15-min TMT exposure as previously reported(6, 24, 67, 89). Furthermore, the alcohol-experienced group demonstrated increased digging and decreased immobility behavior compared to the alcohol-naïve group in male rats (Fig. 7A-D), similar to our previously published work(24). In females, while there were main effects of alcohol-experience on immobility and digging behavior, post-hoc analyses did not yield significant differences at specific timepoints. Interestingly, a history of alcohol experience shifted behavioral response during the TMT exposure towards the subgroup that displayed the greatest interoceptive sensitivity to alcohol (TMT-2) in Fig. 6. This suggests that a history of alcohol exposure may sensitize the stress response to TMT exposure such that its persistent effects result in greater potentiation to the interoceptive effects of alcohol.

**Figure 7.**
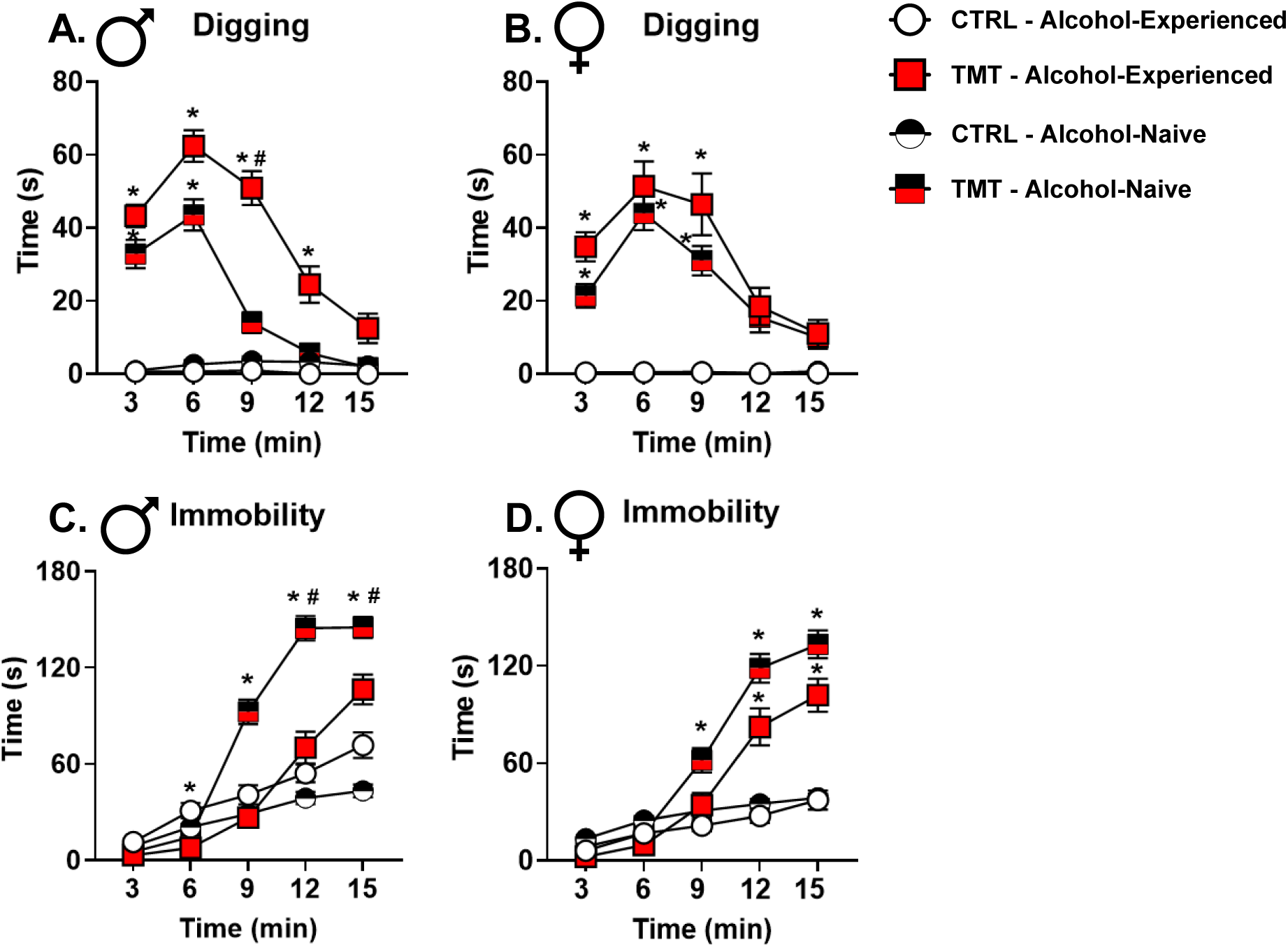
**TMT exposure increases digging and immobility behavior in male and female rats: comparing stress-reactivity in alcohol-experienced and alcohol-naïve cohorts**. During the 15-min TMT exposure, rats engaged in both digging (pushing bedding with the forepaws) and immobility (captures freezing behavior) behaviors. (A) In males rats, the TMT group spent more time digging bedding (F(1, 164)=166.0, p<0.0001) during early time bins compared to the control group in both the alcohol-experienced and alcohol-naïve rats. In the TMT group, alcohol-experienced rats engaged in more digging behavior (F(1, 164)=15.80, p=0.0001) than the alcohol-naïve rats at time bin 9. (B) In female rats, the TMT group spent more time digging bedding (F(1, 150)=130.5, p<0.0001) during early time bins compared to the control group in both the alcohol-experienced and alcohol-naïve rats. There were no differences between the alcohol-experienced and alcohol-naïve groups. (C) In male rats, the TMT group spent more time immobile (F(1, 160)=51.50, p<0.0001) than the control group in the alcohol-naïve group, but not in the alcohol-experienced group. In the TMT groups, the alcohol-experienced rats spent less time immobile (F(1, 160)=10.31, p=0.0016) at time bins 12 and 15 compared to the alcohol-naïve rats. (D) In female rats, the TMT groups spent more time immobile (F(1, 150)=69.14, p<0.0001) compared to the control groups for both alcohol-experienced and alcohol-naïve rats. There was a main effect of alcohol-experience (F(1, 150)=13.75, p=0.0003), but post-hoc tests showed no differences between the alcohol-experienced and the alcohol naïve rats. CTRL/Male/Alcohol-Experienced n=27, TMT/Male/Alcohol-Experienced n=43, CTRL/Male/Alcohol-naïve n=50, TMT/Male/Alcohol-naïve n=48, CTRL/Female/Alcohol-Experienced n=28, TMT/Female/Alcohol-Experienced n=28, CTRL/Female/Alcohol-naïve n=49, TMT/Female/Alcohol-naïve n=49. *p ≤ 0.05 for TMT vs. control groups. ^#^p ≤ 0.05 for alcohol-experienced vs. alcohol-naïve groups.

## DISCUSSION

Exposure to the predator odor TMT potentiated the interoceptive effects of alcohol in male rats through GABA_A_ receptor adaptations in the prelimbic cortex (PrL). The functional, molecular, and behavioral effects of TMT suggest this enhancement may result from a shift in the balance between stimulatory and sedative effects of alcohol to favor greater stimulatory effects. Because the stimulatory effects are associated with the rewarding properties of alcohol, and the sedative effects are associated with aversive properties(90), these adaptations may be one mechanism by which traumatic stress increases alcohol consumption and/or vulnerability to develop AUD, particularly in males. Because men may drink more for reward vs. relief relative to women(91–94), these findings may have identified a novel vulnerability factor by which males escalate drinking following traumatic stress.

Two weeks following TMT exposure, *Gabra1* expression in the PrL was decreased in males, but increased in females. TMT exposure also increased PrL c-Fos expression in males, but not females. Because the PrL is predominantly composed of excitatory, glutamatergic neurons(95), we suspect increased activity of these excitatory neurons, which could be driven by the observed *Gabra1* decrease in the PrL. Previous studies also show that stress is associated with increased glutamatergic neuronal activity in the PrL(48). Further, PTSD is associated with glutamatergic and GABAergic adaptations in the medial prefrontal cortex (mPFC)(64, 65, 96). The fact that acute alcohol administration blocked the TMT-induced increase of c-Fos expression in males suggests that alcohol use may “normalize” the stress adaptations in the PrL. Additionally, TMT exposure attenuated the motor- and startle response-suppressing effects of alcohol in males. These effects led to the hypothesis that TMT exposure may be changing sensitivity to the interoceptive effects of alcohol.

Alcohol produces biphasic effects, starting with stimulation during the ascending limb of the blood alcohol concentration (BAC) curve, followed by sedation during the descending limb of the BAC curve(90, 97). Both stimulation and sedation are components of the interoceptive stimulus of alcohol(31, 32). GABA_A_ receptor activation or NMDA receptor antagonism substitute for the interoceptive effects of alcohol, meaning that the interoceptive effects produced by these drugs are similar to that produced by alcohol, with the degree of substitution depending on the alcohol training dose. At lower alcohol doses (< 1.5 g/kg), GABA_A_ receptor activation achieves full substitution, while NMDA receptor antagonism often results in partial substitution; at higher alcohol doses, this trend reverses with greater recruitment of NMDA receptors(30, 31). Lower alcohol doses tend to be more stimulating and higher alcohol doses tend to be more sedative(41, 42, 98), recruiting GABA_A_ and NMDA receptor contributions, respectively. Thus, we used receptor-specific drugs and site-specific microinjections to dissect the pharmacological and anatomical components of the stressor-induced potentiation to the interoceptive stimulus of alcohol.

Male, but not female, rats showed potentiated sensitivity to the interoceptive effects of 0.5 and 1.0 g/kg alcohol two weeks after TMT exposure. Despite potentiation at low alcohol doses and changes in PrL *Gabra1* expression, TMT exposure did not potentiate the alcohol-like effects of systemically administered pentobarbital. In contrast, we did observe greater alcohol-like effects of the GABA_A_ receptor agonist muscimol microinjected directly into the PrL, but not the anterior insular cortex (aIC), in males after TMT exposure. The discrepancy between systemic pentobarbital vs. site-specific muscimol may reflect differences in drug pharmacology (pentobarbital is an allosteric modulator and muscimol is an orthosteric agonist)(99), and points to specific recruitment of PrL GABA_A_ receptors. These findings demonstrate that GABA_A_ receptors in the PrL are especially important for the increased sensitivity to alcohol following TMT. Interestingly, for both intra-PrL muscimol and MK-801 administration, an increase in locomotor rate was observed following TMT exposure, suggesting that neurons in the PrL are sensitized to the locomotor effects of muscimol and MK-801, both of which contribute to the interoceptive effects of alcohol.

Multiple different findings converge on the hypothesis that TMT exposure results in a shift towards enhanced stimulatory effects of alcohol (i.e., stimulation-sedation balance shifts towards greater stimulation) in males. These include (1) potentiated interoceptive sensitivity to alcohol at low alcohol doses (0.5 and 1.0 g/kg), (2) increased sensitivity to the alcohol-like effects of GABA_A_ activation (muscimol) in the PrL, (3) increased locomotion following muscimol and MK-801 injected into the PrL, (4) decreased *Gabra1* and increased c-Fos expression in the PrL, and (5) blunted sensitivity to the locomotor- and startle response-suppressing effects of alcohol. Because the stimulatory effects of alcohol are associated with its rewarding effects, this shift in the interoceptive effects of alcohol in males may contribute to increased alcohol consumption. However, our previous work did not find increases in operant alcohol self-administration in males following TMT exposure; increases were found in female rats(6). This may be due to several factors including the alcohol consumption measure (binge-like intake or home cage drinking was not assessed) or the differences in alcohol history (the present drug discrimination-trained rats were exposed to considerably more alcohol than the previously reported operant self-administration group), which we describe in previous work(24). Conversely, it is also possible that potentiated sensitivity to the effects of alcohol “protects” against increases in subsequent drinking in male rats.

While female rats did not show changes in interoceptive effects of alcohol, there were notable effects of TMT exposure in females. Females showed an increase in *Gabra1* in the PrL following TMT exposure and diminished sensitivity to the locomotor-enhancing and startle response-suppressing effects of alcohol. Female rats trained on the Pavlovian drug discrimination showed an alcohol dose-dependent decrease in locomotion in the control, but not the TMT group. However, control, alcohol-naïve rats showed increased locomotion following 2 g/kg alcohol that was not observed in the TMT group. This discrepancy may be due to differences in alcohol exposure history and/or the assessments (novel open field vs. a familiar operant chamber). As such, while female rats did not show potentiated interoceptive sensitivity to alcohol, they did show changes in response to alcohol and related neurobiological adaptations.

The reason that females did not show potentiated interoceptive sensitivity to alcohol like males could be due to sex differences in the GABA_A_ receptor(100, 101), which drives the effect of TMT in males. Females showed an opposing effect of TMT on *Gabra1* expression in the PrL relative to males, indicating that females adapt to TMT exposure differently than males. Most importantly, females showed greater alcohol-like effects of systemic pentobarbital than males.

Therefore, females were more sensitive to the GABAA-mediated effects of alcohol, and as such may have been “protected” from the consequences of TMT exposure on increased interoceptive sensitivity to alcohol. This may also account for why alcohol-naïve female rats showed potentiated locomotion when male rats showed attenuated locomotion to the same alcohol dose on the open field test. However, it is important to note that females suffer from PTSD at greater rates than males(1), and female rats are particularly sensitive to the effects of TMT exposure on driving escalations in alcohol self-administration(6). Therefore, this model or methodology may not capture the complex relationship between traumatic stress and interoceptive sensitivity to alcohol in female rats.

Females compared to males with AUD are more likely to drink in response to negative emotions and have co-occurring psychiatric comorbidities(91–93). In people with PTSD and alcohol dependence, both men and women drank alcohol for coping motives, but only men showed greater drinking for reward enhancement motives(94). As such, women may be more motivated to drink alcohol to relieve negative emotionality, whereas men may be more likely to drink for reward. This may also suggest that our previous work showing escalations in alcohol self-administration in female, but not male rats after the same TMT exposure protocol(6) may reflect drinking to relieve a negative state, rather than reward-related drinking. Therefore, this work may have identified a sex-specific, novel vulnerability factor by which traumatic stress affects vulnerability for AUD.

Upon examination of individual differences in stress-reactive behaviors during the TMT exposure, we found that high stress-reactive male rats (higher D/I ratios; TMT-2 subgroup) showed the greatest increase in sensitivity to 1.0 g/kg alcohol. This suggests that there is individual variability in response to traumatic stress that is associated with changes in interoceptive sensitivity to alcohol. This finding is congruent with the clinical observation that only a minority of people who experience traumatic stress develop PTSD or co-morbid AUD(1). Interestingly, a history of alcohol exposure shifted the behavioral response during TMT to favor greater stress reactivity in males, consistent with our previous work(24). Therefore, a history of alcohol exposure prior to a traumatic stress event may sensitize the stress response, influencing the likelihood that the traumatic stressor will alter sensitivity to alcohol.

The present study has limitations. Firstly, we were not able to examine individual differences via the TMT subgrouping for the PrL and aIC microinjection substitution experiments given the smaller sample sizes. Additionally, the different stress-reactive profiles during TMT exposure between alcohol-experienced and alcohol-naïve rats suggest that the persistent effects of TMT exposure may be different between these conditions. Finally, while the changes in the alcohol-like effects of muscimol injected into the PrL indicate a direct role of GABA_A_ receptors in stressor-induced potentiation of interoceptive sensitivity to alcohol, this effect was accompanied by increased locomotion during the test session. This suggests that the effects may not be entirely specific to the interoceptive effects of alcohol. However, intra-PrL MK-801 also increased locomotion and substitution was not altered, making such an explanation less tenable and providing more evidence that TMT exposure enhances the stimulatory effects of alcohol.

Together, these data show that TMT exposure enhanced interoceptive sensitivity to alcohol through GABA_A_ receptor adaptations in the PrL in male, but not female rats. Additional behavioral and molecular evidence suggested greater stimulatory effects of alcohol in males.

Moreover, alcohol history enhanced stress reactivity behavior during the stressor, which was linked to greater potentiation of the interoceptive effects of alcohol. These findings may provide insights into the high comorbidity between PTSD and AUD by elucidating how traumatic stress affects alcohol sensitivity.

## Acknowledgements

This work was supported in part by the National Institutes of Health [AA026537 and AA011605 (JB)] and by the Bowles Center for Alcohol Studies. RET was supported in part by NS007431 and AA029946.

## Author Contributions (CRediT)

**Ryan E. Tyler**: Conceptualization, Methodology, Formal analysis, Investigation, Data Curation, Writing – Original Draft, Writing – Review and Editing, Visualization, Supervision, Funding Acquisition. **Maya N. Bluitt**: Formal analysis, investigation. **Kalynn J. Van Voorhies**: Formal analysis, investigation. **Wen Liu**: Formal analysis, investigation. **Sarah N. Magee**: Investigation. **Elisabeth R. Pitrolo**: Investigation. **Victoria L. Cordero**: Investigation. **Laura C. Ornelas**: Methodology, Investigation. **Caroline G. Krieman**: Investigation. **Brooke N. Bender**: Formal analysis, Investigation. **Alejandro M. Mosera**: Investigation. **Joyce Besheer**: Conceptualization, Resources, Writing – Review and Editing, Supervision, Project administration, Funding acquisition.

