## Supplemental Materials for "The persistent effects of predator odor stressor enhance interoceptive sensitivity to alcohol through GABA_A_ receptor adaptations in the prelimbic cortex in male, but not female rats"

### SUPPLEMENTAL METHODS AND MATERIALS

#### *Animals*

Male and female Long-Evans rats (Envigo) arrived at 7 weeks of age and were housed individually under a 12-hour light/dark cycle, with experiments conducted during the light phase. Rats were handled daily for 1 minute for 1 week before experiments began. For Pavlovian drug discrimination (Exp. 4) and alcohol metabolism (Exp. 5), rats were fed daily to maintain 85% of free-feeding weight. Alcohol-naïve rats (Exp. 1-3) had ad libitum access to food. All rats had ad libitum access to water. Department of Comparative Medicine staff at UNC-Chapel Hill provided continuous care, and all procedures followed NIH guidelines for animal care and use.

#### *Predator odor stressor: TMT exposure*

Rats were exposed to 2,5-dihydro-2,4,5-trimethylthiazoline (TMT) or water for controls in separate test chambers (45.72 × 17.78 × 21.59 cm; UNC Instrument Shop, Chapel Hill, NC). Experimental set-up were identical to those used in previous work(6, 24, 68, 86; see 67 for detailed protocol description). Briefly, rats were transported from the vivarium in the home cage to a separate, well-ventilated room that contained the test chambers in which rats were exposed to TMT. Only one rat was placed in each chamber. The length of the back wall of the test chambers was opaque white with two opaque black side walls and a clear, plexiglass front wall to enable video recordings and a clear sliding lid. The bottom of the chamber was covered with a layer of white bedding, also used in our previous publications(6, 24, 68, 86). A small, metal basket was hung on the right-side wall (17.8 cm above the floor) to hold a piece of filter paper. 10 μL of TMT (2,5-dihydro-2,4,5-trimethylthiazoline) or 10 uL of water for controls was pipetted onto the filter paper in the metal basket immediately prior to putting the rat in the chamber. The control group was always run before the TMT group to prevent odor contamination. The odor exposure session lasted 15 mins (except for a 10-min exposure in Exp. 5 - alcohol metabolism experiment) and was video recorded for evaluation of behavior using ANY-maze Video Tracking System (Version 6.12, Stoelting Co. Wood Dale, IL). After TMT exposure, rats were returned to the home cage and remained undisturbed in the home cage for 2 weeks as an "incubation period" prior to starting behavioral testing or sacrifice for molecular assessments.

***Experiment 1: Gene expression after TMT exposure***

Male and female alcohol-naïve rats underwent TMT exposure and were sacrificed 2 weeks later. Brains were collected for gene expression analyses in the PrL and aIC using qRT-PCR analyses as previously described(24, 75, 86). Both NMDA and $GABA_A$ receptor subunit genes were examined (see Table 1 for primer sequences). Male and female rats were counterbalanced within the same cohort, enabling direct statistical comparison between sex. Tissue collection and gene expression analysis methods were identical to previously published work(24, 75, 86) and are described briefly below.

*Brain tissue collection*

Rats were anesthetized with isoflurane, and brains were collected and flash frozen with isopentane (Sigma-Aldrich, MI). Brains were stored at -80°C prior to brain region sectioning. Brains were sectioned on a cryostat until the region of interest was visible and then tissue for the region of interest was excised with a micro-punch tool.

*RNA extraction* – The RNeasy Mini Kit (Qiagen; Venlo, Netherlands) was used to extract RNA from brain tissue according to the manufacturer's instructions. A Spectrophotometer (Nanodrop 2000, ThermoScientific) was used to determine RNA concentration and purity for each sample.

*Reverse Transcription* – The SuperScript™ III First-Strand Synthesis System (ThermoFisher Scientific) was used to reverse transcribe RNA into cDNA according to the manufacturer's instructions. Next, all samples were diluted 1:5 with water and stored at −20 °C before RT-PCR experiments.

*RT-PCR* – All experiments used the QuantStudio3 PCR machine (ThermoFisher). All samples were run in duplicate with a 96-well plate, using 10 μL total volume per well with the following components: PowerUp Syber green dye (ThermoFisher, containing ROX dye for passive reference), forward and reverse primers (Eton Biosciences Inc.; North Carolina, United States), and cDNA template. The PCR was run with an initial activation for 10 min at 95 °C, followed by 40 cycles of the following: denaturation (95 °C for 15 s), annealing/extension (60 °C for 60 s). Melt curves were run for all genes to verify synthesis of only one amplicon. *Gapdh* served as the housekeeper gene. Supplemental Table 1 displays all primer sequences.

***Experiment 2: Neuronal sensitivity to alcohol and TMT exposure***

Alcohol-naïve male and female rats underwent the TMT exposure. Two weeks later, rats received a single oral gavage of alcohol (2.0 g/kg, i.g.) or water (i.g.) for vehicle controls prior to being sacrificed and perfused (90-min pretreatment time). Brains were collected and c-Fos immunoreactivity via immunohistochemistry was quantified in the PrL and aIC. Males and females were counterbalanced within the same cohort, enabling direct comparison of sex, but sex was not used as a biological variable in analyses.

*Brain tissue collection and sectioning*

90-min after the alcohol or water administration, rats were deeply anesthetized with isoflurane. Rats were transcardially perfused with 0.1 M PBS (4°C, pH=7.4) followed by 4% paraformaldehyde (PFA, 4°C, pH=7.4). Brains were extracted and stored in PFA for 24 h at 4°C.

Brains were then rinsed with 0.1 M PBS and moved to a 30% sucrose in 0.1 M PBS solution. Forty µm coronal sections were collected on a freezing microtome. Sections were stored in cryoprotectant at -20 °C until starting immunohistochemistry (IHC).

*c-Fos immunohistochemistry*

Coronal sections (free floating) were rinsed in PBS (0.1 M) before a 5-min wash in 1% hydrogen peroxide. Next, sections were blocked in 3% normal goat serum (NGS; Vector Labs, Burlingame, CA) in 0.3% Triton X-100 for 2 h. Sections were then incubated in rabbit anti-c-Fos antibody (1:4000 in 3% NGS + 0.1% Triton; Synaptic Systems, Gottingen, Germany; Lot# 226003/3–45) for 16 h at 4°C. This was followed by an incubation with biotinylated goat anti-rabbit secondary antibody (1:200 in 3% NGS + 0.1% Triton X-100; Vector Labs, Burlingame, CA; Lot # 140893). Next, sections were incubated in Vectastain Elite ABC HRP (Vector labs, Burlingame, CA). Lastly, brain sections were treated with diaminobenzidine (Sigma-Aldrich, St. Louis, MO) and then mounted on glass slides.

Brain section images were obtained using the Olympus CX41 light microscope (Olympus America, Center Valley, PA). The analyses were conducted with Image-Pro Premier image analysis software (Media Cybernetics, Rockville, MD). Bilateral immunoreactivity data (c-Fos-positive cells/mm2) were taken from a minimum of 2 sections/brain region/animal (e.g., 4 data points per animal), conducted by an experimenter blind to animal groups. These data were then averaged to obtain one value for each subject. The regions examined were the aIC and the PrL (bregma +3.00 to +3.72, see Fig. 2E for areas of quantification). Brain regions of quantification have been described previously(5, 18, 20, 24).

***Experiment 3: Behavioral sensitivity to alcohol and TMT exposure***

Alcohol-naïve male and female rats underwent TMT exposure. Two weeks later, rats received alcohol (2.0 g/kg, i.g.) or water (i.g.) for vehicle controls. 20-mins later, rats were placed in a novel open-field for 20 min to assess locomotion. The following day, rats were tested on acoustic startle response 20-min after alcohol (2 g/kg, i.g.) or water (i.g.) for vehicle controls. Alcohol vs. water pre-treatment was counterbalanced across the open field and acoustic startle response testing days. Males and females were run in separate cohorts and were analyzed separately.

*Open-Field Test*

Rats were transported to the test room 30 min before the experiment began for acclimation. Rats received alcohol (2 g/kg, i.g.) or water (eq. v/v) 20 min prior to the start of the open field test (20 min test duration). Rats were placed in the center of the open-field arena (43 cm × 43 cm; Med Associates, St. Albans, VT) and locomotor activity was recorded by 32 orthogonal infrared beams using Activity Monitor software (Med Associates). Lights were not turned on inside the open field test chamber or during acclimation to the test room, creating a dark environment for this test. Red light was used for the experimenter.

*Acoustic Startle Response Test*

24-hours after the open field test, rats were evaluated on an acoustic startle response test using an acoustic startle response system (S-R Lab; San Diego Instruments, San Diego, CA) after administration of alcohol (2 g/kg, i.g., 20 min pretreatment). Rats were transported in their home cages to the behavioral testing room and allowed to acclimate for 30 min prior to testing. Rats were placed in a cylinder-shaped, plexiglas animal enclosure located within a sound-attenuating test chamber that included an exhaust fan, a sound source, and an internal light that was turned off during the test. At the start of each test, rats underwent a 5-min habituation period during which 60 dB of background white noise was present. The background noise was present during the entire test session. The test session consisted of 30 trials of a 100 ms burst of a 110 dB startle noise identical to previously published work from the lab(6). Each trial was separated by a 30- to 45-s randomized intertrial interval. Startle response (amplitude) was measured with a high-accuracy accelerometer mounted under the animal enclosure and analyzed with SR-Lab. This test was conducted under normal light conditions.

***Experiment 4: Interoceptive sensitivity to alcohol following TMT exposure***

Male and female rats underwent Pavlovian drug discrimination training to discriminate alcohol (2.0 g/kg, i.g., 20-min pretreatment) from water (i.g.) as we have previously described(17, 18, 24, 25, 35, 37). Once the discrimination was acquired, rats underwent TMT exposure and remained undisturbed in the home cage. 2 weeks after TMT exposure, rats began testing on an alcohol dose response curve (0.0, 0.5, 1.0, 2.0 g/kg, i.g.) across different days (i.e., one dose tested per day in ascending order – 2.0, 1.0, 0.5 and 0.0 g/kg). A subset of rats underwent substitution testing of $GABA_A$ receptor agonists and a NMDA receptor antagonist following the alcohol dose-response testing days. Substitution experiments included both systemic injections (pentobarbital, 5 mg/kg, i.p.) in males and females, as well as site-specific injections to both the prelimbic cortex (PrL, bilateral coordinates: AP +3.2, ML ±0.6 mm, DV −2.0 mm) or the anterior insular cortex (aIC, bilateral coordinates: AP +2.5, ML ±4.0 mm, DV −4.6 mm) in males only. Males and females were trained and tested as separate cohorts and analyzed separately.

See below for detailed description.

*Apparatus* – On one side of the chamber, cue lights were located on either side of a liquid receptacle. A photobeam detector was used to measure head entries into the liquid receptacle.

*Sucrose Access Training* – Training started with three 50-min sessions which provided sucrose (26 %) randomly throughout the session. The probability of sucrose presentation decreased from the first to the last session and by the last 10 min of the final session rats received approximately 0.75 sucrose presentations/min.

*Acquisition Training* – Training sessions were conducted 5 days per week (M-F). On these training sessions, alcohol (2 g/kg, IG) or water (IG) was administered and the rats were immediately placed in the chambers. After a 20-min delay (to allow for blood and brain alcohol levels to rise), the 15-min training session began. During the alcohol training session, the offset of each 15-sec cue light presentation (10 random presentations throughout the session) was followed by sucrose presentation (26%, 0.1 mL, 4 sec) into the liquid receptacle. In contrast, during water training sessions, sucrose was not delivered following the offset of the cue light presentations. Training sessions occurred on a double alternating schedule (A, A, W, W…). Testing started following at least 12 alcohol and water training sessions. Using the mean discrimination score from the previous 2 water and 2 alcohol sessions, the alcohol session score – water session score had to be ≥ 2 to meet criteria for testing. Only rats that met criteria for discrimination were included in the analyses.

*Pavlovian drug discrimination: testing*

Test sessions consisted of the standard 20-min timeout period, and then a single 15-s cue light presentation. Alcohol was administered in 4 separate doses (0.0, 0.5, 1.0, and 2.0 g/kg, i.g.) on separate testing days per dose. No sucrose was delivered following the offset of the light presentation.

*Systemic pentobarbital substitution*

Rats were administered pentobarbital (5 mg/kg, i.p.) using a 10 mg/mL solution in saline and injecting 0.5 mL/kg body weight. Immediately following pentobarbital administration rats received oral administration of water identical to water training sessions. Rats were then placed in the operant box and underwent a test session (single light presentation following a 20-min delay, no sucrose delivery).

*Surgeries*

Following training on the alcohol drug discrimination task, a subset of male rats underwent surgeries to implant bilateral cannulae targeting either the prelimbic cortex (PrL) or the anterior insular cortex (aIC). Rats were anesthetized with isoflurane (5% for induction and 2% for maintenance), then received bilateral implantation of 26-gauge guide cannulae (Plastics One, Roanoke, VA) aimed to terminate 2 mm above the anterior insular cortex (aIC, bilateral coordinates: AP +2.5 mm, ML ±4.0 mm, DV −4.6 mm) or the prelimbic region of the PFC (PrL, bilateral coordinates: AP +3.2 mm, ML ±0.6 mm, DV −2.0 mm). Coordinates were based on(82). For analgesia, banamine (2.5 mg/kg) was administered subcutaneously the day prior to surgery, the day of surgery, and the day following surgery. Cannulae placements for successful hits and rats included in the analyses are depicted in Fig. 5C (PrL) and Fig. 5D (aIC). Following 1-week recovery period, all rats received 4 additional training sessions (2 Alcohol, 2 Water) before TMT exposure.

*Microinjections*

All drugs for microinjections were delivered through injectors extending 2 mm below the guide cannulae. Site-specific microinjections were delivered using a microinfusion pump (Harvard Apparatus, MA) through 5.0 µl Hamilton syringes connected to 33-gauge injectors (Plastics One, VA). The rate of injection for all microinjections was 0.5 µL/min. Muscimol was injected at a volume of 0.5 µl/side over 1 min. MK-801 was injected at a volume of 1 µL/side over 2 min. aIC: 10 mM muscimol 0.5 uL per side, PrL: 2.5 mM muscimol 0.5 uL per side. PrL: 2.0 mg/mL

MK801 1 uL per side. MK-801 was not tested in the aIC. The injectors remained in place for another 2-min after the infusion to allow for diffusion.

*Perfusions and cannulae verification*

Rats were transcardially perfused identical to Experiment 2 (c-Fos Experiment) to confirm correct cannula placement. Brain tissue was stained with cresyl violet to verify cannulae placement. Only data from rats with cannulae/injector tracts determined to be in the target brain regions were used in analyses. 5 rats were removed from analyses of the PrL due to incorrect cannulae placement. 1 rat were removed from analyses of the aIC due to incorrect cannulae placement.

*Drugs*

Alcohol (95% w/v) was diluted in tap water to a concentration of 20% (v/v). Sodium Pentobarbital (Patterson Veterinary, MA) was dissolved in saline (0.9%) to a concentration of 10 mg/mL. Muscimol (R&D systems, Minneapolis, MN) was dissolved in saline (0.9%) to produce a 10mM (for aIC microinjections) and a 2.5 mM (for PrL microinjections) solution. MK-801 (Tocris Biosciences, Cat. # 0924) was dissolved in saline to a concentration of 2.0 mg/mL.

***Experiment 5 – Alcohol Metabolism***

Male and female rats underwent a 10-min TMT exposure (or water exposure for controls). Two weeks later, all rats received a 1.0 g/kg oral gavage of alcohol. Tail blood was collected at intervals of 20-, 60-, 90-, and 120-minutes post-gavage to assess blood alcohol concentration (BAC, mg/dL). Sex was used as a biological variable. Blood samples were centrifuged for 1 min at 10 rcf, and 10-30 µL of plasma were collected from each sample. Collected plasma was evaluated by an Analox AM1 Analyzer (200 mg/dL standard) to determine BAC at each time interval.

**Supplemental Data Analysis**

For Experiment 1 (gene expression experiment, Fig. 1), the ΔΔCt method was used to calculate fold change relative to controls. Fold changes were normalized such that the average male control (used as the ultimate control group) fold change was equal to 1. A 2-way ANOVA was run with group (CTRL and TMT) and sex (male and female) as the between subjects factors. If a significant interaction effect was observed, post-hoc Bonferroni's multiple comparisons test was run to evaluate the effects of TMT exposure in males and females separately. The initial sample sizes were n=10/group/sex. Some samples were lost from the original sample size due to experimental error (RNA extraction errors or poor sample quality). This resulted in a lower sample size in the PrL relative to the aIC.

For Experiment 2 (c-Fos immunoreactivity experiment, Fig. 2), the average number of c-Fos positive cells per $mm^2$ were used as the dependent measure. A 2-way ANOVA was run with group (CTRL and TMT) and pre-treatment condition (alcohol or water) as the between subjects factors. If a significant interaction effect was observed, post-hoc Bonferroni's multiple comparisons test was run to evaluate the effects of TMT exposure in water (vehicle)- and alcohol-treated groups separately. The initial sample size was n=8/group/alcohol pretreatment for both males and females. Some samples were not usable for analyses due to experimental error or tissue damage in the area of quantification.

For Experiment 3 (open field and acoustic startle response, Fig. 3), the total distance traveled in the open field during the total 20-min session was the primary dependent measure for locomotion. For ASR, the average of the peak startle response for the 30-trials was used. A 2-way ANOVA with group (TMT vs. CTRL) and pretreatment condition (water or alcohol) as the between subjects factors was run. We *a priori* hypothesized that alcohol (2 g/kg) would affect these behavioral measures. Therefore, planned comparisons were conducted if there was a main effect of alcohol to determine effects of alcohol administration in the CTRL and TMT groups separately. The initial sample size was n=12/group/alcohol pretreatment for both males and females. Some rats were lost to analysis due to poor intragastric injection.

For Experiment 4 (drug discrimination experiment, Fig. 4-6), the primary dependent measure was the discrimination score. The discrimination score was calculated as [number of head entries into the liquid receptacle during the 15-s light presentation] minus [number of head entries during the 15-s period immediately *before* the light presentation]. This measure serves as a readout for the interoceptive effects of the training drug (2.0 g/kg alcohol, i.g.)(25). Locomotor rate data was also collected during the drug discrimination testing as the number of beam breaks per min. For drug discrimination and locomotor rate variables, a 2-way ANOVA with group as a between subjects factor and alcohol dose as a within subjects factor was used. If a group x dose interaction effect was observed, post-hoc Bonferroni multiple comparisons were conducted to test group differences (CTRL vs. TMT) at each alcohol dose tested. For pentobarbital substitution experiments, a 2-tailed, unpaired t-test was conducted between CTRL and TMT groups. For site-specific substitution experiments (Fig. 5), a 1-tailed, unpaired t-test with Welch's correction was used. 1-tailed test was used because the effect of potentiated sensitivity was already established, and a Welch's correction was used due to differences in sample size between the CTRL and TMT groups. Only rats that met criteria for discrimination both prior to TMT exposure and after TMT exposure were included. Criteria for discrimination was defined as [alcohol discrimination score – water discrimination score ≥ 2]. Sample sizes were as follows: CTRL/male: n=21, TMT/male: n=40, CTRL/female: n=18, TMT/female: n=15. A subset of these rats were tested for pentobarbital substitution: Male/CTRL n=15, Male/TMT n=19,

Female/CTRL n=9, Female/TMT n=9. A subset of male rats were used for site-specific microinjections: PrL/Muscimol/MK-801: CTRL n=6, TMT=16. aIC/Muscimol: CTRL n=6, TMT n=7. MK-801 was not tested in the aIC. The same rats for PrL cannulae were used to test both muscimol and MK-801 effects. Some rats were lost to analyses due to poor intragastric gavage or cannulae misplacement.

Substitution analysis for the 2.0 g/kg alcohol training dose was assessed by using the average training dose discrimination score prior to TMT exposure (shown as a dotted line on graphs), and then using a 1-sample t-test to compare the pre-TMT alcohol training dose average ("hypothetical" value) to the discrimination scores following substitution drug (e.g., pentobarbital) administration. If there was a statistically significant difference between the substitution discrimination scores relative to the pre-TMT training dose average, this was considered "no substitution" for the training dose. If there was not a statistically significant difference, this was considered "substitution" for the training dose. Substitution analysis was not conducted for site-specific microinjection experiments due to the low sample size.

For the sub-group analyses of Exp. 4 (Fig. 6), the purpose was to segregate the male rats exposed to TMT into high and low stress-reactive subgroups as an exploratory investigation of individual differences in response to the TMT exposure. The behaviors quantified were digging and immobility and are identical to our previous work(6, 63, 76). Digging was defined as pushing bedding with the forepaws. This behavior was usually observed as rats pushing bedding towards the TMT source to create a mound or barrier between themselves and the TMT source. Immobility was defined as the absence of movement other than respiration for 2 seconds or longer. While this measure does capture inactivity, it mainly captures freezing behavior especially given that the TMT chambers are novel and only 15-min in duration. The ratio of time spent digging over time spent immobile (D/I) during the TMT exposure was used to sub-group rats into high (D/I > 1, TMT-2) and low (D/I <1, TMT-1) stress reactivity similar to our previous work(6, 63). The TMT-2 group engaged in more digging behavior relative to immobility while the TMT-1 group engaged in more immobility behavior relative to digging as defined by the experimenter subgroups. Sample sizes that met criteria for alcohol drug discrimination analyses are as follows: CTRL: n=21, TMT-1: n=27, and TMT-2: n=13. The post-TMT alcohol-dose response and pentobarbital substitution data was then re-analyzed using these 3 groups. A 1-way ANOVA with group as the between subjects factor was run. Sidak's multiple comparisons test was used to determine group differences at each alcohol dose tested for the alcohol dose response. For pentobarbital substitution and the 1 g/kg alcohol dose analysis, if there was a main effect of group, Tukey's multiple comparisons test was used to compare differences between the three groups (CTRL, TMT-1 and TMT-2).

The behavioral response during the 15-min TMT exposure for males and females was measured for all experiments and is displayed in Fig. 7. This resulted in 2 groups with distinct alcohol-exposure history: i.e., the alcohol-naïve rats (Exp. 1, 2, 3) and the alcohol-experienced rats (Exp. 4). Sample sizes are as follows: For males, CTRL/alcohol-naïve: n=50, TMT/alcohol-naïve: n=48, CTRL/alcohol-experienced: n=27, and TMT/alcohol-experienced: n=43. For females, CTRL/alcohol-naïve: n=49, TMT/alcohol-naïve: n=49, CTRL/alcohol-experienced: n=28, and TMT/alcohol-experienced: n=28. Time spent digging and immobile were assessed overtime in 3-min bins (3, 6, 9, 12, and 15-min for 5-time bins). A 3-way ANOVA was run with alcohol-experience and TMT exposure as between-subjects factors, and timepoint as the within subjects factor. Post-hoc tests were conducted if a significant TMT exposure x alcohol-experience x time-point effect was observed. Tukey's multiple comparisons test was used for post-hoc tests. Differences between the TMT vs. CTRL and the alcohol-naïve vs. alcohol-experienced groups at each timepoint separately were reported.

For Experiment 5 (alcohol metabolism, Supplemental Fig. 1), blood alcohol concentration was analyzed using a 3-way ANOVA with TMT exposure as a between subjects factor, sex as a between subjects factor, and timepoint as a within-subjects factor. Post-hoc tests were not run because there were no interaction effects. The sample sizes was as follows: CTRL/male: n=4, TMT/male: n=8, CTRL/female: n=4, TMT/female: n=8.

**SUPPLEMENTAL RESULTS**

*TMT exposure sex-dependently altered Gabra1 expression in PrL and upregulated NMDA receptor subunits in aIC in males (**Figure 1**).*

*Prelimbic Cortex*

For all NMDA receptor subunits (Fig. 1A-D) tested, there was no effect of TMT exposure or sex on gene expression. For *Gabra1* (Fig. 1E), there was a significant effect of sex ($F(1, 28)=9.52$, $p=0.005$), no effect of TMT, and a significant TMT x sex interaction ($F(1, 28)=18.12$, $p=0.0002$). Interestingly, TMT exposure decreased *Gabra1* in males ($p=0.01$), but increased *Gabra1* in females ($p=0.01$). For all other GABA-A subunits (Fig. 1F-I) tested, no main effect of TMT, sex nor interaction was found.

*Anterior Insular Cortex*

*Grin1* and *Grin2a* (Fig. 1J-K) expression was not affected by TMT exposure or sex. *Grin2b* (Fig. 1L) did not show a main effect of TMT or sex but there was a significant TMT exposure x sex interaction ($F(1, 33)=5.10$, $p=0.03$), with *Grin2b* upregulated as an effect of TMT exposure in males ($p=0.03$), but not in females. Likewise, *Grin2c* (Fig. 1M) did not show a significant effect of TMT or sex but did show a significant TMT x sex interaction ($F(1, 33)=6.23$, $p=0.018$), with

TMT exposure increasing expression in males (p=0.04) but not in females. *Gabra1* and *Gabra4* were not affected by TMT exposure or sex (Fig. 1N, O). Some $GABA_A$ subunits showed no main effect of TMT or sex but did show significant TMT x sex interactions (*Gabarb2* (Fig. 1P): F(1, 33)=3.97, p=0.05; *Gabrd* (Fig. 1Q): F(1, 33)=5.21, p=0.03; *Gabrg2* (Fig. 1R): F(1, 31)=5.04, p=0.03). Post-hoc tests did not yield statistically significant effects of TMT or sex.

***Figure 2*** *- TMT exposure increased PrL c-Fos expression in males, which is not observed in rats pre-treated with alcohol*

In males in the PrL (Fig. 2A), there was a main effect of TMT exposure (F(1, 22)=12.70, p=0.002), no main effect of alcohol dose, but a significant TMT exposure x alcohol dose interaction (F(1, 22)=6.83, p=0.02). Post-hoc tests showed that there was a greater number of c-Fos positive cells in the TMT group compared to controls in the water-treated group (p=0.0003); however, this effect was not observed in the alcohol-treated group. PrL c-Fos expression in females (Fig. 2B) did not show an effect of TMT exposure or alcohol pretreatment, nor an interaction effect. c-Fos expression in males in the aIC (Fig. 2C) did not show a main effect of TMT exposure or alcohol dose, nor an interaction effect. aIC c-Fos expression in females (Fig. 2D) did not show a main effect of TMT exposure, sex, nor interaction effect.

***Figure 3*** *– Alcohol produced locomotor and startle response effects in the control group, but not in the TMT-exposed group*

For the open-field test in males (Fig. 3A), a main effect of alcohol was observed on total distance traveled (F(1, 39)=10.63, p=0.002). There was no main effect of TMT exposure or interaction. Planned comparisons showed that alcohol decreased locomotion in males in the control group (t(18)=4.31, p=0.0004) but not in the TMT group (p=0.14). For the open-field test in females (Fig. 3B), there was a main effect of alcohol on total distance traveled (F(1, 41)=4.64, p=0.04), but no effects of TMT exposure or interaction. Planned comparisons showed that alcohol increased locomotion in the control group ($t(20)=2.08$, $p=0.05$), but not in the TMT group ($p = 0.34$).

For startle response in males (Fig. 3C), alcohol induced an overall decrease in average peak startle response as supported by a significant main effect of alcohol ($F(1, 40)= 1.65$, $p=0.002$), and no effect of TMT exposure or interaction. Planned comparisons show that alcohol decreased average peak startle response in the control group ($t(19)=2.79$, $p=0.01$), but not in the TMT group, although there was a trend for a decrease ($t(21)=1.93$, $p=0.07$). For startle response in females (Fig. 3D), alcohol induced an overall decrease in average peak startle response as supported by a significant main effect of alcohol ($F(1, 37)=5.35$, $p=0.03$), and no effect of TMT exposure or interaction. Planned comparisons showed that alcohol decreased average peak startle response in the control group ($t(18)=2.18$, $p=0.04$), but not in the TMT group ($p=0.34$).

***Figure 4*** *– TMT exposure potentiated interoceptive sensitivity to alcohol in male but not female rats.*

For male rat discrimination prior to TMT exposure (Fig. 4A), there was a main effect of alcohol on the discrimination scores ($F(1, 59)=262.2$, $p<0.0001$), confirming appropriate discrimination between the effects of alcohol from water. There was no effect of group prior to TMT exposure. For male locomotion prior to TMT exposure (Fig. 4B), there was no effect of group or alcohol dose. For female rat discrimination prior to TMT exposure (Fig. 4C), there was a main effect of alcohol on discrimination score ($F(1, 31)=211.8$, $p<0.0001$), again confirming appropriate discrimination between the effects of alcohol from water. There was no effect of group on discrimination score prior to TMT exposure. For female rat locomotion prior to TMT exposure (Fig. 4D), there was a main effect of alcohol ($F(1, 31)=6.48$, $p=0.02$), but no effect of group or interaction. These data confirm discrimination of alcohol from water and show that there were no group differences prior to the TMT exposure.

Two weeks after TMT exposure, testing was conducted on a full alcohol dose response curve. Male rats (Fig. 4E) showed a main effect of alcohol dose ($F(2.38, 140.3)=63.83$, $p<0.0001$), a main effect of TMT exposure ($F(1, 59)=11.32$, $p=0.001$) and an alcohol dose x TMT exposure interaction ($F(3, 177)=3.09$, $p=0.03$). Post-hoc tests showed higher discrimination scores at the 0.5 g/kg dose ($p=0.02$) and the 1.0 g/kg dose ($p=0.009$) in the TMT group vs. the control group, but not at 0.0 or 2.0 g/kg doses. For locomotion in males after TMT exposure (Fig. 4F), there was no effect of alcohol dose or TMT exposure. Discrimination in female rats (Fig. 4G), showed a main effect of alcohol dose ($F(3, 93)=27.03$, $p<0.0001$), but no effect of TMT exposure or interaction. For locomotion in female rats (Fig. 4H), there was a main effect of alcohol dose ($F(3, 93)=11.77$, $p<0.0001$), no main effect of TMT exposure, but a significant alcohol dose x TMT exposure interaction ($F(3, 93)=3.70$, $p=0.01$). Post-hoc tests comparing differences in locomotor rate between each alcohol dose and vehicle (0.0 g/kg) showed that in the control group, locomotion was decreased at every alcohol dose compared to vehicle ($p<0.05$). However, there were no differences in locomotion between alcohol doses and vehicle in the TMT group. The dotted lines on panels E and F represent the group mean (groups combined because this was prior to the TMT exposure) of the alcohol session prior to the TMT exposure taken from panel A and C, receptively, and serve as a comparative for full substitution. For males, the mean was $6.41 \pm 0.42$ SEM, and for females the mean was $6.27 \pm 0.23$ SEM.

Pentobarbital (5 mg/kg, i.p.) substitution in male rats did not show a difference between the TMT and control groups on discrimination score (Fig. 4I), but locomotor rate (Fig. 4J) was lower in the TMT group compared to controls ($t(32)=2.45$, $p=0.02$). Comparing both the control and TMT group discrimination scores to the group average alcohol discrimination score prior to TMT exposure, showed that there was no substitution for the alcohol training dose (CTRL: $t(14)=4.82$, $p = 0.0003$; TMT: $t(18)=4.33$, $p = 0.0004$). Pentobarbital substitution in female rats did not show a group difference for discrimination score (Fig. 4K) or locomotor rate (Fig. 4L). However, unlike in males rats, pentobarbital produced substitution for the alcohol training dose (pentobarbital discrimination scores relative to the average discrimination score prior to TMT exposure - CTRL: ($t(8)=1.61$, $p=0.14$, TMT: $t(8)=0.41$, $p=0.69$). Similar to the alcohol dose response curves, a dotted line on the discrimination score graphs (panel I and K) reflect the mean of the pre-TMT training dose discrimination scores.

***Figure 5*** *– TMT exposure potentiated the alcohol-like stimulus effects of* $GABA_A$ *receptor agonism in the prelimbic cortex in male rats*

TMT exposure increased the alcohol-like stimulus effects of muscimol injected into the PrL (Fig. 5B, $t(11.19)=1.82$, $p=0.047$). TMT exposure increased the locomotor rate following muscimol injection in the PrL (Fig. 5C, $t(13.91)=2.86$, $p=0.0064$). There was no effect of TMT exposure on the alcohol-like stimulus effects of MK-801 injected into the PrL (Fig. 5D). TMT exposure increased the locomotor rate following PrL microinjection of MK-801 (Fig. 5E, $t(8.06)=5.86$, $p=0.0002$). Muscimol injected into the aIC did not result in group differences for (Fig. 5G) discrimination scores or (Fig. 5H) locomotor rate. Sample sizes were insufficient for substitution analyses.

***Figure 6*** *– Heightened behavioral response to TMT exposure is associated with higher interoceptive sensitivity to alcohol, but not with the alcohol-like effects of pentobarbital.*

The D/I ratio was higher in the TMT group compared to controls (Fig. 6A, $t(65)=2.27$, $p=0.03$). For the 3 groups, there was a main effect of group for D/I ratio (Fig. 6B, $F(2, 64)=13.74$, p=0.0003). By definition of the subgroups, the D/I ratio was higher in TMT-2 compared to TMT-1 ($p<0.0001$) and controls ($p<0.0001$). For interoceptive sensitivity to alcohol, there was a main effect of group ($F(2, 58)=10.20$, $p=0.0002$), alcohol dose ($F(2.429, 140.9)=74.30$, $p<0.0001$), and a group x alcohol dose interaction effect ($F(6, 174)=2.904$, $p=0.01$). Post-hoc comparisons showed significant differences between the CTRL and TMT-2 group at 1.0 g/kg alcohol. When analyzing the 1.0 g/kg alcohol dose separately, there was a main effect of group ($F(2, 58)=8.047$, $p=0.0008$). Post hoc comparisons showed higher discrimination scores in the TMT-2 group compared to both the CTRL and TMT-1 groups, but no difference between TMT-1 and CTRL groups.

***Figure** 7 – TMT exposure increased digging and immobility behavior in male and female rats: comparing alcohol-experienced and alcohol-naïve cohorts*

For digging in males (Fig. 7A), there was a main effect of TMT exposure ($F(1, 164)=166.0$, $p<0.0001$), a main effect of alcohol-experience ($F(1, 164)=15.80$, $p=0.0001$), and a main effect of time ($F(3.323, 544.9)=64.01$, $p<0.0001$). There was a significant interaction effect for TMT exposure and alcohol-experience ($F(1, 164)=24.32$, $p<0.0001$), TMT exposure and time ($F(4, 656)=63.08$, $p<0.0001$), and alcohol-experience and time ($F(4, 656)=4.99$, $p=0.0006$). There also an interaction effect between TMT exposure x alcohol-experience x time ($F(4, 656)=6.056$, $p<0.0001$). Post-hoc tests in alcohol-naïve rats showed greater digging behavior in the TMT group compared to controls at time bins 3 and 6, but not 9, 12, and 15. In alcohol-experienced rats, the TMT group engaged in more digging behavior compared to controls at time bins 3, 6, 9, and 12, but not 15. In the TMT group, alcohol-experienced rats engaged in more digging compared to alcohol-naïve rats at time bin 9, but not 3, 6, 12, and 15. In the control group, there were no differences in digging behavior at any time-bin between the alcohol-experienced and alcohol-naïve groups. For digging behavior in female rats (Fig. 7B), there was a main effect of TMT exposure ($F(1, 150)=130.5$, $p<0.0001$), no main effect of alcohol-experience, and a main effect of time ($F(3.08, 461.8)=34.29$, $p<0.0001$). There was a TMT exposure x time interaction effect ($F(4, 600)=33.66$, $p<0.0001$), but no TMT exposure x alcohol-experience or alcohol-experience x time effect. There was no TMT exposure x alcohol-experience x time interaction effect. In alcohol-naïve rats, the TMT group engaged in more digging behavior compared to controls at time bins 3, 6, and 9, but not 12 and 15. Likewise in alcohol-experienced rats, the TMT group engaged in more digging behavior compared to the CTRL group at time bins 3, 6, and 9, but not 12 and 15. Post-hoc tests between alcohol-experienced and alcohol-naïve rats were not conducted because there was no main effect of alcohol experience or alcohol-experience x TMT exposure interaction effect. For immobility behavior in male rats (Fig. 7C), there was a main effect of TMT exposure ($F(1, 160)=51.50$, $p<0.0001$), a main effect of alcohol-experience ($F(1, 160)=10.31$, $p=0.0016$), and a main effect of time ($F(2.89, 461.6)=237.7$, $p<0.0001$). There was a significant interaction effect for TMT exposure and alcohol-experience ($F(1, 160)=47.68$, $p<0.0001$), TMT exposure and time ($F(4, 640)=63.80$, $p<0.0001$), and alcohol-experience and time ($F(4, 640)=10.30$, $p<0.0001$). There was a significant TMT exposure x alcohol-experience x time interaction effect ($F(4, 640)=16.23$, $p<0.0001$). In alcohol-naïve rats, the TMT group engaged in more immobility behavior compared to controls at time bins 9, 12, and 15, but not 3 and 6. In alcohol-experienced rats, the TMT group engaged in less immobility compared to controls at time bin 6, but there were no differences at time bin 3, 9, 12, and 15. In the TMT group, alcohol-naïve rats engaged in more immobility behavior compared to alcohol-experienced rats at time bins 9 and 12, but not 3, 6, and 15. In the control group, there were no difference in immobility behavior between alcohol-experienced and alcohol-naïve rats. For immobility in female rats (Fig. 7D), there was a main effect of TMT exposure ($F(1, 150)=69.14$, $p<0.0001$), alcohol-experience ($F(1, 150)=13.75$, $p=0.0003$), and time ($F(2.674, 401.1)=155.2$, $p<0.0001$). There was an interaction effect for TMT exposure and alcohol-experience ($F(1, 150)=3.90$, $p=0.05$), TMT exposure and time ($F(4, 600)=70.12$, $p<0.0001$), but not for alcohol-experience and time. There was a significant interaction effect of TMT exposure x alcohol-experience x time ($F(4, 600)=2.483$, $p=0.04$). In alcohol-naïve rats, the TMT group engaged in more immobility behavior compared to the control group at time bins 9, 12, and 15, but not at 3 and 6. In alcohol-experienced rats, the TMT group engaged in more immobility behavior compared to the control group at time bins 12 and 15, but not 3, 6, and 9. In both the control and TMT groups, there were no differences in immobility behavior between the alcohol-experienced and alcohol-naïve rats.

**SUPPLEMENTAL TABLES**

***Table S1*** – Primer Sequences

| Gene Name | Forward Primer (5'-3') | Reverse Primer (5'-3') |
|---|---|---|
| *Gabra1* | GAGCACGCAGAGTCCATGA | TTCTTCATCACGGGCTTGTCC |
| *Gabra4* | ACCGTGTACTTTCACCTCAGAC | GTGAGGACTGTGGTTATTCCAAAT |
| *Gabrb2* | TGGGGTGCTTTGTCTTTGTCT | TCATGTGGGTCCATCTTGTTGA |
| *Gabrd* | TTTTATACAGCATCCGCATCACC | TAGCTCTCCAGGTCCAGCAT |
| *Gabrg2* | GAAGTCTCTGCCCAAGGTCTC | AATGGTAGGGGCAGGGTTTT |
| *Grin1* | CTATGACAACAAGCGCGGAC | GCCCGTCATGTTCAGCATTG |
| *Grin2a* | GCGGGAACCCGCTAAACC | GCAATACCAGCAAGGTCCAGT |
| *Grin2b* | GGGTCACGCAAAACCCTTTC | CCTTGTTTTTGACGCCCCTG |
| *Grin2c* | CAACGTCTTGGTTCCCCTCA | CTTGGGCTTCTCCTCTCAGC |
| *Gapdh* | AACGACCCCTTCATTGAC | TCCACGACATACTCAGCAC |

Table S1 shows the forward and reverse primer sequences for all gene targets in the gene expression experiment (Exp. 1, Fig. 1). *Gapdh* was used as the housekeeper gene.

**SUPPLEMENTAL FIGURES**

***Figure S1* – No effect of TMT exposure or sex on alcohol metabolism.**

These experiments were conducted to assess the effects of TMT exposure and sex on alcohol metabolism, as changes in alcohol levels at the time of testing (20 min post alcohol administration) could influence discrimination behavior results independently of neurobiological adaptations. There was no main effect of TMT exposure or sex on alcohol metabolism. As expected, there was a significant main effect of timepoint on blood ethanol concentrations ($F(3, 60)=17.53$, $p<0.001$), with no interaction between TMT exposure, timepoint, and sex. These findings suggest that the effects of TMT exposure and sex differences were likely not due to variations in alcohol metabolism.

**Supplemental Fig. 1**

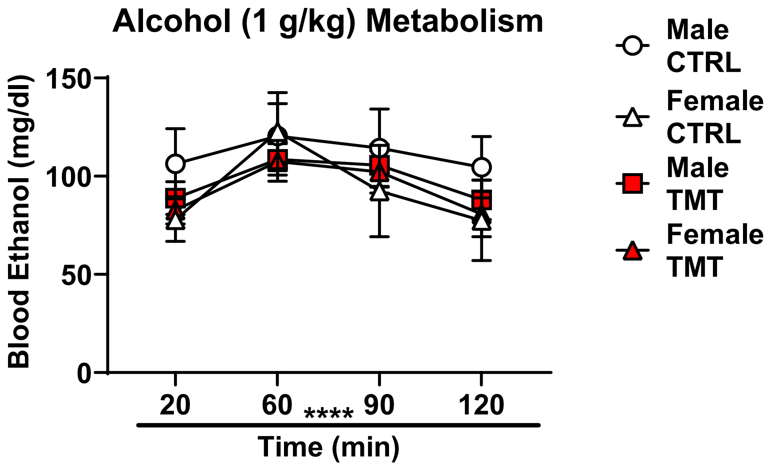


**Figure S1** – There was a main effect of time-point ($F(3, 60)=17.53$, $p<0.0001$) on ethanol levels in the blood. There was no effect of TMT exposure or sex on ethanol levels over time. CTRL/male: n=4, TMT/male: n=8, CTRL/female: n=4, TMT/female: n=8. ****$p<0.0001$.
